# Broadly-recognized, cross-reactive SARS-CoV-2 CD4 T cell epitopes are highly conserved across human coronaviruses and presented by common HLA alleles

**DOI:** 10.1101/2022.01.20.477107

**Authors:** Aniuska Becerra-Artiles, J. Mauricio Calvo-Calle, Marydawn Co, Padma P. Nanaware, John Cruz, Grant C. Weaver, Liying Lu, Catherine Forconi, Robert W. Finberg, Ann M. Moormann, Lawrence J. Stern

**Author notes:** These authors contributed equally (Aniuska Becerra-Artiles, J. Mauricio Calvo-Calle).

## Abstract

Sequence homology between SARS-CoV-2 and common-cold human coronaviruses (HCoVs) raises the possibility that memory responses to prior HCoV infection can impact the T cell response in COVID-19. We studied T cells recognizing SARS-CoV-2 and HCoVs in convalescent COVID-19 donors, and identified a highly conserved SARS-CoV-2 sequence S_811-831_, with two overlapping epitopes presented by common MHC-II proteins HLA-DQ5 and HLA-DP4. These epitopes were recognized by CD4^+^ T cells from convalescent COVID-19 donors, mRNA vaccine recipients, and by low-abundance CD4^+^ T cells in uninfected donors. TCR sequencing revealed a diverse repertoire with public TCRs. CD4^+^ T cell cross-reactivity was driven by the remarkably strong conservation of T cell contact residues in both HLA-DQ5 and HLA-DP4 binding frames, with distinct patterns of HCoV cross-reactivity explained by MHC-II binding preferences and substitutions at secondary TCR contact sites. These data highlight S_811-831_ as a highly-conserved CD4^+^ T cell epitope broadly recognized across human populations.

## Introduction

SARS-CoV-2 infections result in a wide spectrum of clinical presentation from asymptomatic to severe disease and death, with most cases resulting in relatively mild symptoms (Wiersinga et al., 2020). However, in an important fraction of the population that includes older individuals or in individuals with comorbidities, infections frequently develop into a severe disease with high mortality (Gerges Harb et al., 2020). Age-related variations in the innate and adaptive host response to SARS-CoV-2 (Koch et al., 2021) and genetic polymorphisms (Soveg et al., 2021; Wickenhagen et al., 2021) play a critical role in the disparity of the clinical outcome. Deficient anti-viral immunity in nasal epithelial cells (Ziegler et al., 2021), reduced type I interferon (Vanderbeke et al., 2021; Ward et al., 2021), atypical peripheral blood cytokine profile (Mazzoni et al., 2021), recruitment and activation of neutrophils are important in the local and systemic pathogenesis (Koch et al., 2021; Vanderbeke et al., 2021). Against this background, the role of T cells in severe disease is unsettled (reviewed in (Sette and Crotty, 2021)). In severe cases, there are dysregulated T cell responses (Rydyznski Moderbacher et al., 2020), and potentially pathogenic tissue-resident CD4^+^ T cells that express IL-17A and GM-CSF are found in the lungs of SARS-CoV-2 patients (Zhao et al., 2021). In support of a central role played by T cells in protection against SARS-CoV-2, studies have shown that low CD4^+^ and CD8^+^ T cell counts are associated with severe disease (Chen et al., 2020; Du et al., 2020), and that peak severity is inversely correlated with the frequency of SARS-CoV-2-specific IFN-γ-producing CD8^+^ T cells (Rydyznski Moderbacher et al., 2020). In addition, early CD4^+^ T cell responses are associated with mild disease (Peng et al., 2020; Tan et al., 2021a).

Six other coronaviruses, in addition to the now pandemic SARS-CoV-2, are known to infect humans: the highly-pathogenic SARS and MERS beta-coronaviruses, which caused constrained outbreaks initially in 2002 and 2012, respectively, and the much less pathogenic alpha-coronavirues 229E and NL63, and betacoronaviruses OC43 and HKU1, which are believed to have emerged within the last few centuries, and now circulate seasonally and are associated with mild common-cold symptoms (Cui et al., 2019; Forni et al., 2017; Gaunt et al., 2010). Shortly after the discovery of the original SARS virus, it was shown that T cells from unexposed individuals could recognize naturally processed and presented SARS antigens even before infection (Chen et al., 2005; Gioia et al., 2005; Yang et al., 2009). Serological cross-reactivity and sequence homology in major proteins between SARS and the circulating common-cold coronaviruses (HCoVs) led to the suggestion that previous infections with HCoVs might have elicited the cross-reactive SARS-specific responses in unexposed individuals (Meyer et al., 2014). A limited number of studies argued against this hypothesis (Chen et al., 2005; Yang et al., 2009). T cells from individuals not previously infected with SARS that recognized a SARS-derived CD8+ T cell epitope had an impaired response when compared with CD8+ T cells from previously SARS-infected individuals, and did not recognize a peptide covering the homologous region of the 229E coronavirus (Chen et al., 2005). In another study, of SARS-specific immune responses in uninfected donors, the CD4+ T cell response was limited to CD45RA+ T cells that were considered naïve (Yang et al., 2009). However, the first study was limited to just one of the four circulating coronaviruses and the selected peptide has low sequence homology. In addition, it is now known that CD4+ effector memory T cells responding to viral antigens can re-express CD45RA (Di Mitri et al., 2011), and these cells are implicated in protective responses to pathogens (Tian et al., 2017).

Immune cross-reactivity among the human coronaviruses has received much recent attention with the spread of SARS-CoV-2 in human populations, because of the possibility that the varied spectrum of COVID-19 disease potentially could be explained in part by pre-existing immunity elicited by prior exposure to circulating seasonal coronaviruses (reviewed in (Bonifacius et al., 2021; Lipsitch et al., 2020). In support of protective responses, both unexposed and SARS-CoV-2-infected individuals have cross-reactive antibodies to both SARS-CoV-2 and HCoV spike protein, which are boosted by SARS-CoV-2 infection (Ng et al., 2020; Shrock et al., 2020), and some of these antibodies can neutralize SARS-CoV-2 in vitro (Ng et al., 2020). Additionally, individuals with a recent HCoV infection have been reported to have less-severe COVID-19 (Sagar et al., 2021). However, a similar study did not find that previous infection with HCoVs reduced COVID-19 severity (Gombar et al., 2021). The relatively short lifetime of the serological response (Edridge et al., 2020) used in these studies to assess prior exposure may complicate this approach to identification of cross-reactive responses between SARS-CoV-2 and HCoVs. In general, cellular responses to coronavirus infections are longer-lived than serological responses (Bilich et al., 2021; Le Bert et al., 2020; Liu et al., 2006). Several studies have observed T cell responses to SARS-CoV-2 antigens in uninfected donors (reviewed in (Grifoni et al., 2021)). Up to 50% of people who had not been exposed to SARS-CoV-2 had significant T cell reactivity (Braun et al., 2020; Grifoni et al., 2020; Le Bert et al., 2020; Weiskopf et al., 2020). However, SARS-CoV-2 epitopes recognized in uninfected donors represent only a small fraction of the overall response observed after SARS-CoV-2 infection (Le Bert et al., 2020; Tarke et al., 2021; Weiskopf et al., 2020). Some studies report that SARS-CoV-2 responsive T cells in uninfected donors have low avidity (Bacher et al., 2020; Dykema et al., 2021; Saini et al., 2021), although another study reported comparable responses for some epitopes (Mateus et al., 2020). Recent evidence for a potentially protective role of pre-existing SARS-CoV-2-specific T cells in COVID-19 comes from a study of people who had been exposed to SARS-CoV-2 but did not test positive for infection (i.e. undergoing abortive infections), who have early T cell responses to SARS-CoV-2 replication complex proteins that are highly conserved coronaviruses and may have helped to prevent productive infection (Swadling et al., 2021). Prexisting memory T cells respond more quickly to spike mRNA vaccine than do newly-elicted T cells, with levels that correlate with antibodies specific for SARS-CoV-2 spike protein, suggesting a possible supportive role for cross-reactive T cells in COVID-19 vaccination (Loyal et al., 2021).

Despite the high prevalence of HCoV infection and substantial sequence homology with SARS-CoV-2 (Chen et al., 2021), the relationship between pre-infection SARS-CoV-2-specific T cell reponses and prior HCoV infection is not clear. Mateus et al. expanded SARS-CoV-2 epitope specific T cell lines from non-infected donors, and tested their reactivity against the corresponding HCoV peptides (Mateus et al., 2020). Of 42 T cell lines recognizing SARS-CoV-2 cross-reactive peptides, only 9 recognized the homologus HCoV epitopes (Mateus et al., 2020). Subsequent bioinformatic analysis of pre-existing T cell responses to SARS-CoV-2 suggests that prior cross-reactivity with HCoVs can explain only a fraction of these (Tan et al., 2021b). It is possible that SARS-CoV-2-specific responses observed before infection might have been primed by other stimuli besides homologous HCoVs, and in fact one report demonstrated that a CD4^+^ T cell epitope in SARS-CoV-2 spike protein cross-reacts with naturally processed Bacteroidales and Klebsiella antigens (Lu et al., 2021). However Dykema et al., were able to validate cross-reactive recognition of both SARS-CoV-2 and NL63 homologs for all five TCRs tested (Dykema et al., 2021).

In order to help clarify the role of cross-reactive T cell responses in COVID-19, we investigated SARS-CoV-2 spike protein responses targeted by cross-reactive T cells isolated from previously uninfected donors and convalescent COVID-19 individuals. We used an unbiased screen to identify epitopes targeted by these cells. We systematically screened T cell lines cross-reactive between SARS-CoV-2 spike protein epitopes and corresponding HCoV antigens that were raised from previously uninfected donors and from convalescent COVID-19 donors. We identified a highly-conserved immunodominant peptide broadly recognized in most donors by a polyclonal and polyfunctional CD4^+^ T cell response. Two epitopes within this sequence are presented by HLA molecules common in populations worldwide. T cells that recognize this peptide also respond to the corresponding HCoV epitopes with similar avidity. The response is characterized by a broad repertoire of TCRs, with subsets responding to different patterns of HCoV variations. The conserved sequence S_811-831_ (KPSKRSFIEDLLFNKVTLADA) can be used to follow cross-reactive responses during SARS-CoV-2 infection or vaccination, and might be a good candidate for inclusion in pan-coronavirus vaccination strategies.

## Results

### T cell responses to coronavirus antigens in COVID-19 and previously uninfected donors

To begin to characterize the cross-reactive T cell response to SARS-CoV-2 and HCoVs, we measured responses to overlapping peptide pools covering the spike (S) proteins of the four HCoVs, and the spike (S), membrane (M), nucleoprotein (N), and envelope (E) proteins of SARS-CoV-2, using blood samples from recovered COVID-19 donors at convalescence and a set of previously uninfected donors that include both unexposed pre-pandemic donors sampled 2015-2018, and seronegative asymptomatic individuals sampled contemporaneously with the COVID-19 donors. We measured IFN-γ secretion in response to antigenic stimulation directly ex-vivo, and also after a single stimulation in-vitro with HCoV S peptide pools (S pools) in order to expand cross-reactive T cell populations. Representative IFN-γ ELISpot ex-vivo data for a COVID-19 donor and a pre-pandemic donor are shown in Fig. 1A, and representative in-vitro expanded data are shown in Fig. 1B. Summaries of the ex-vivo responses for 12 COVID-19 and 20 uninfected donors are shown in Fig. 1C, and summaries of in-vitro expanded cell responses for 7 COVID-19 and 12 uninfected donors are shown in Fig. 1D.

**Figure 1.**
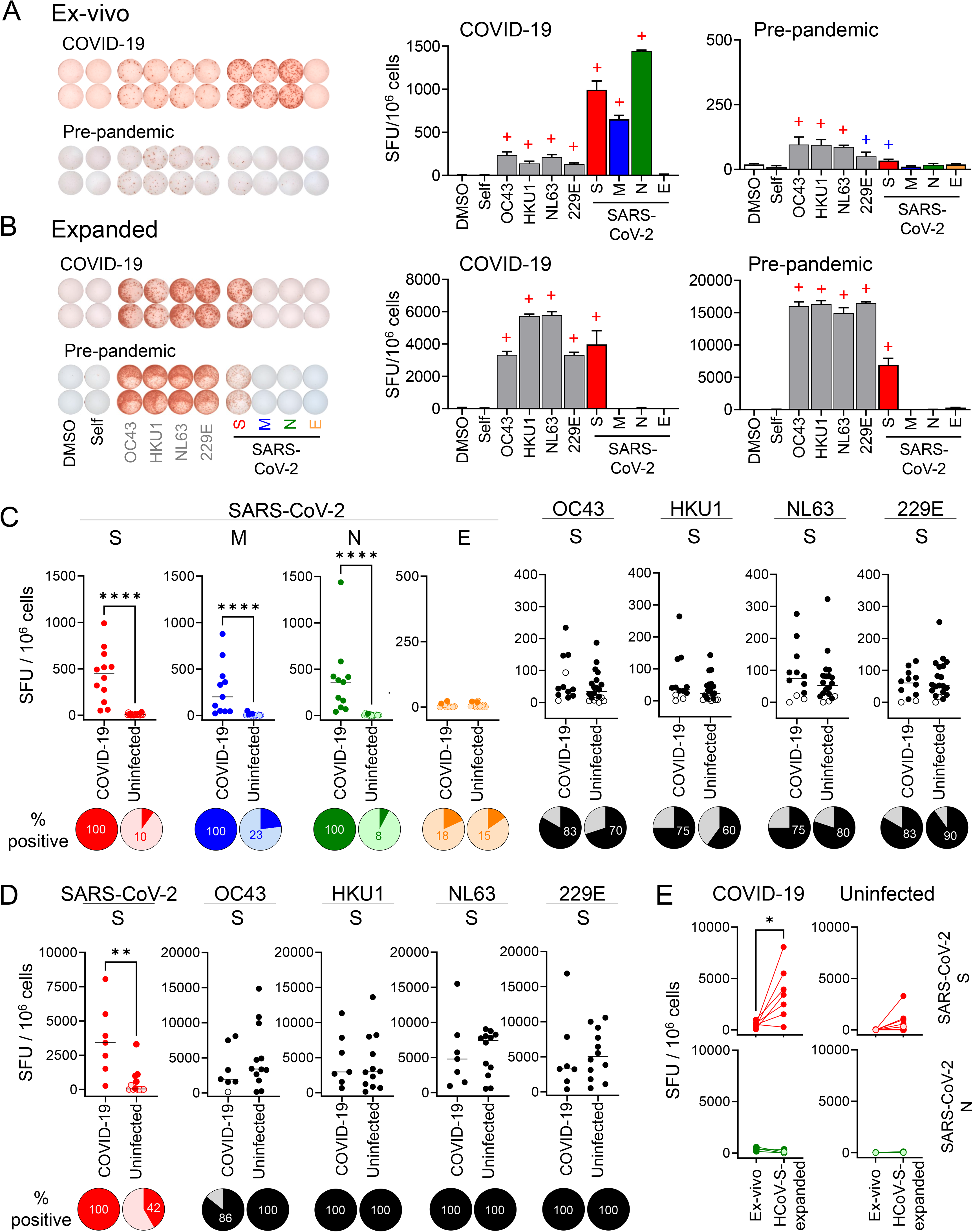
Responses to coronavirus antigens in COVID-19 and uninfected donors. Responses to S pools from coronaviruses OC43, HKU1, NL63, and 229E (gray), and of S (red), M (blue), N (green) and E (orange) pools from SARS-CoV-2 were studied in COVID-19 donors at convalescence and uninfected donors. Representative ex-vivo responses in one COVID-19 donor at convalescence and one uninfected pre-pandemic donor (A) and responses to re-stimulation with the indicated antigens after in-vitro expansion with HCoV S pools in same donors (B). IFN-γ ELISpot images (left) and average (± standard deviation) of replicate wells (bar graphs) are presented; “+” signs above bars represent positive responses by DFR1X (blue) or DFR2X (red) tests (see methods). Summary of ex-vivo responses in 12 COVID-19 donors at convalescence and 20 uninfected donors (C) and summary of responses of in-vitro expanded T cells in 7 COVID-19 at convalescence and 12 uninfected donors (D); statistical analysis by Mann-Whitney test (significant responses: **** p<0.001, ** p<0.01); pies: percentage of positive responses (dark color) for each group/condition. E. Comparison of the responses to re-stimulation with SARS-CoV-2 S or N peptides pools, before and after expansion with HCoV S pools, in COVID-19 at convalescence and uninfected donors; statistical analysis by paired t-test: * p=0.021.

A representative COVID-19 donor (d0801, **Supplemental Table S1**) showed strong IFN-γ responses to peptide pools from SARS-CoV-2 S, M, N but not E protein (Fig. 1A). Responses to S pools from the HCoVs were weaker, but clearly distinguishable from self-peptide and vehicle controls. To validate the HCoVs responses, we tested whether responding T cell populations could proliferate in-vitro in after stimulation with S pools from the four HCoVs. Responses to all HCoVs S pools were expanded (~27-fold) by this treatment (Fig. 1B). Off-target expansion appeared to be minimal, as responses to SARS-CoV-2 M, N, or E pools were not expanded. Responses to SARS-CoV-2 S pool also were expanded by stimulation with the HCoVs S pool (~4-fold), indicating that some fraction of the SARS-CoV-2-responsive T cell population was able to cross-react with HCoV homologs.

A pre-pandemic donor (L38) exhibited IFN-γ T cell responses to S pools from each of the four HCoVs, presumably elicited by by prior sesonal exposure to these viruses, and also to SARS-CoV-2 S peptides (Fig. 1A). This donor was sampled in 2016 before emergence of SARS-CoV-2 into humans, and so the response to SARS-CoV-2 S peptide suggests a possible cross-reactive response with T cells elicited by prior HCoVs infection. To test this, we expanded T cells with HCoVs S pools as just described (Fig. 1B). SARS-CoV-2 S-specific responses expanded strongly (~90-fold) after heterologous stimulation with a HCoVs S pool, similarly to the HCoVs-specific responses (~51-fold). This indicates that T cell populations from this unexposed donor responsive to SARS-CoV-2 also are cross-reactive with HCoVs homologs.

Similar responses were observed throughout the entire COVID-19 and uninfected study groups (Fig. 1C-D, **Supplemental Table S1**). Ex-vivo responses to SARS-CoV-2 peptide pools were observed in 100% of COVID-19 donors for the S, M and N proteins (Fig. 1C, filled circles). Only 2 of the 11 donors showed responses to the E protein pool. The responses were relatively strong, especially for the S protein. As previously observed in other studies (Le Bert et al., 2020; Mateus et al., 2020; Nelde et al., 2021; Tan et al., 2021b), responses to these SARS-CoV-2 antigens also were observed in uninfected donors, with both the fraction of donors exhibiting positive responses, as well as the numbers of responding T cells observed for these donors substantially lower than for the COVID-19 donors (Fig. 1C). Responses to the SARS-CoV-2 S protein in uninfected donors were relatively weak (23±18 SFU/106 cells), and some positive responses might have been below the limit of detection in our ex vivo assay. After in-vitro expansion, 42% of uninfected donors had positive IFN-γ responses specific for SARS-CoV-2 S pool (Fig. 1D), compared with 10% assayed directly ex-vivo.

Ex-vivo responses to HCoVs S pools were observed in most COVID-19 and uninfected donors, with all donors responding to at least one of the HCoVs (Fig. 1C). As expected, expansion of HCoVs-S-specific cells was observed for donors from the uninfected individuals, such that after expansion essentially everyone was positive for S pools from each virus (Fig. 1D). No significant differences in the magnitude of the responses or frequency of positive responses between COVID-19 and uninfected donors were observed, or between the different HCoVs.

T cell populations from COVID-19 donors responding to SARS-CoV-2 S peptides expanded after stimulation with HCoVs S pools (average 8-fold increase), indicating that responding T cells could recognize both SARS-CoV-2 and homologous HCoVs epitopes (Fig. 1E, top). The expansion was specific for S-derived antigens, as no expansion of N-specific responses was observed for either COVID-19 or uninfected donors (Fig. 1E, bottom). T cell populations from uninfected donors responded to S pool from SARS-CoV-2, a virus to which they had not been exposed, and these responses also expanded after stimulation (average 66-fold increase) with homologous epitopes. Thus, both uninfected and COVID-19 donors exhibited T cell responses cross-reactive between corresponding HCoVs and SARS-CoV-2 antigens.

### Identification of cross-reactive peptides

To identify epitopes responsible for these cross-reactive T cell responses, we screened overlapping peptide libraries covering the full sequence of the SARS-CoV-2 spike protein. As before, we assessed responses using T cells from three COVID-19 donors at convalescence enriched for cross-reactive populations by expansion in vitro with HCoV S pools. We used a two-step pool-deconvolution approach to identify individual peptides involved in the cross-reactive response. First, responses to peptides grouped into pools of 10 were measured (Fig. 2A), and second, pools exhibiting positive responses were deconvoluted and retested to identify individual peptides (Fig. 2B). In this manner, we identified responses to three pairs of overlapping peptides: 163/164 (S_811-831_, green), recognized in all three donors tested; 190/191 (S_946-966_, pink), recognized in two donors; and 198/199 (S_986-1006_, blue), recognized by a single donor (Fig. 2C).

**Figure 2:**
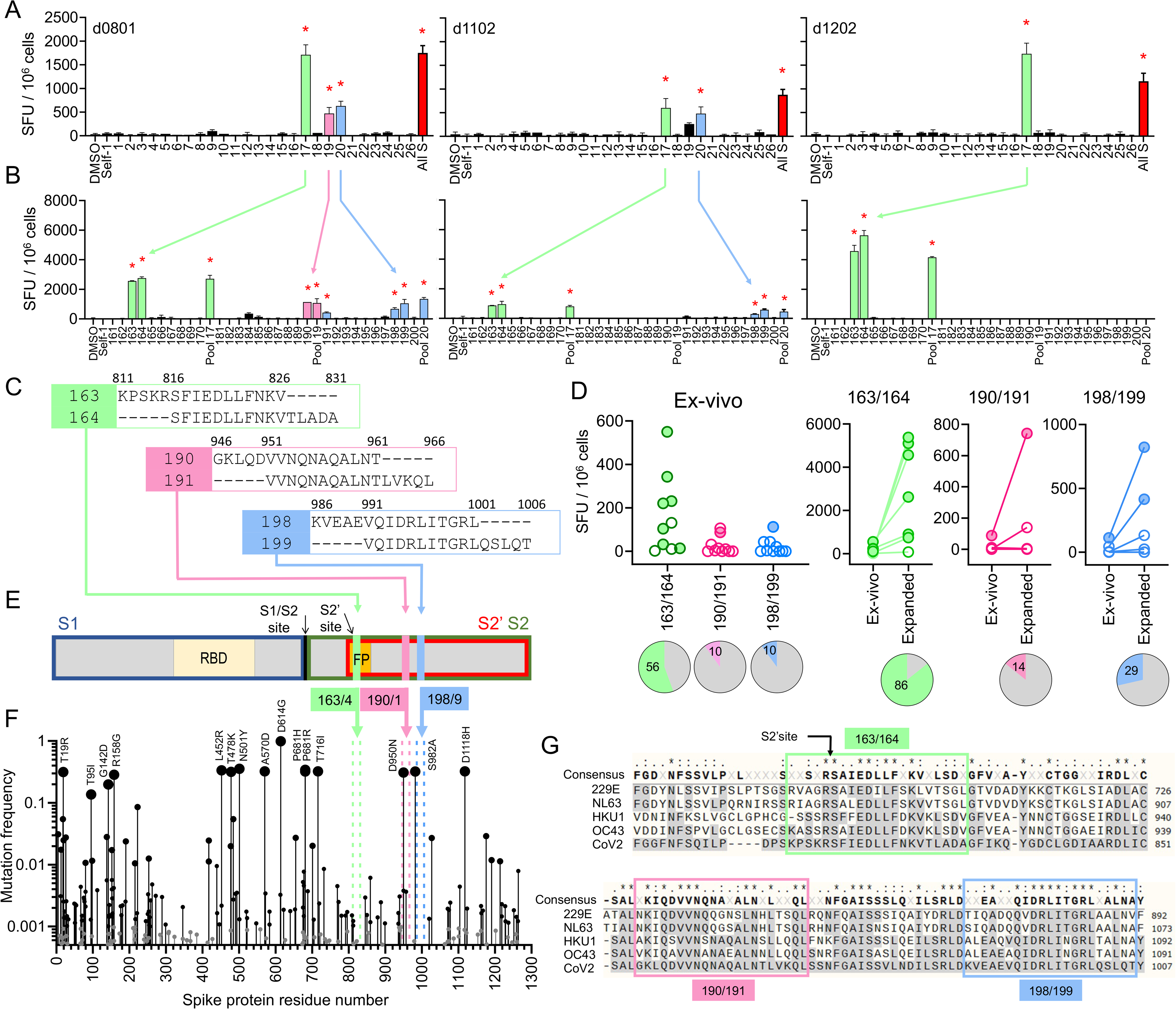
Identification of cross-reactive peptides: A. Responses of in vitro expanded cross-reactive T cells from three COVID-19 donors at convalescence (expanded with HCoV S pools) to re-stimulation with pools of 10 overlapping peptides from SARS-CoV-2 S protein were measured by IFN-γ ELISpot (pool number indicated on X-axis; “All S” is the pool of all S peptides; DMSO and Self-1 are negative controls).. B. Positive pools were deconvoluted to identify responses to individual peptides using ELISpot (peptide number indicated in X-axis; negative controls and parent pool also included) For both, A and B red stars indicate positive responses by DFR2X (Moodie et al., 2012). C. Amino-acid sequences of candidate epitopes identified; sequences of each pair of overlapping peptides and amino acid position in the protein are shown. D. Ex-vivo responses (by IFN-γ ELISpot) to each pair of candidate epitopes identified in B, measured in COVID-19 donors (n= 10); comparison of ex-vivo response to responses of in-vitro expanded cells for the three pairs of peptides; positive responses shown as filled circles and negative responses shown as empty circles. Pies show the percentage of positive responses. E. Schematic of SARS-CoV-2 spike protein. Location of peptides 163/164 (light green), 190/191 (pink), and 198/199 (blue) in the protein is shown; also shown are RBD (receptor binding domain), FP (fusion peptide), cleavage sites (S1/S2 and S2’), and cleavage products S1 (blue box), S2 (green box), and S2’ (red box). F. Mutation frequency in the spike protein from SARS-CoV-2. Size of circles indicate the mutation frequency range. Location of the candidate epitopes in the protein is shown with colored vertical broken lines. Common mutations are also indicated. G. Sequence alignment of S protein from SARS-CoV-2 (bottom) and HCoVs (229E, NL63, HKU1 and OC43) containing the sequences of the candidate epitopes (enclosed in the boxes).

We evaluated the responses to peptides 163/164, 190/191, and 198/199 in additional convalescent COVID-19 donors. In the 10 donors analyzed, 5 showed ex-vivo responses to 163/164, and one donor each recognized 190/191 and 198/199 (Fig. 2D, left). After in-vitro expansion with HCoV S pools, and additional donor was positive for peptide 163/164 (Fig. 2D, right). Thus 163/164 is recognized by a substantial fraction of COVID-19 donors across HLA types (**Supplemental Table S1)**.

All three cross-reactive epitopes identified derive from the S2’ domain of the S protein (Fig. 2E). The 163/164 sequence contains the S2’ cleavage site and the fusion peptide (FP), critical for viral entry (Xia, 2021). 190/191 is in the first heptapeptide repeat region and 198/199 is between heptapeptide repeats 1 and 2. These regions are highly conserved among SARS-CoV-2 variants, including delta (B1.617.2) and omicron (B.1.1.529) variants of concern, with a mutation frequency <0.01 for most positions except S_950_ in 190/191 (Fig. 2F). These regions also are highly conserved when compared the four HCoVs (Fig. 2G), again reflecting their probable functional importance. Overall, these results and observations indicate the 163/164 region is a broadly-recognized immunogenic hotspot in which mutations are highly restricted.

### Functional characteristics of cross-reactive T cell populations

To assess functional characteristics of T cell populations responding to these peptides, we performed a basic phenotypic analysis of the in-vitro-expanded cross-reactive T cells (Fig. 3). Expanded cells had a large CD4+ T cell population (Fig. 3A,B), with a predominantly effector-memory phenotype (Fig. 3C). Intracellular cytokine secretion analysis showed that cells responding to re-stimulation with SARS-CoV-2 S pool (CoV-2 S), or the individual peptides 163 and 164, were exclusively within the CD4+ T cell population, and produced mainly IFN-γ, with some TNF-α, very little IL-2, and mobilized CD107a to surface, suggesting a Th1 population with cytotoxic potential (Fig. 3D). Furthermore, the expanded cells were polyfunctional. Around half of the responding cells produced one or two cytokines along with mobilizing CD107a, though also frequent were cells expressing only CD107a or only IFN-γ (Fig. 3E). These trends were consistent for the 3 donors tested. t-SNE analysis of pooled samples (ctrl, CoV-2 S, 163 and 164) reveals two major clusters (g1-g2) along with some disperse populations (Fig. 3F). Cluster g1 includes cells producing high IFN-γ, TNF-α and mobilizing CD107a. Cluster g2 include cells that still mobilize CD107a but produce less IFN-γ and no TNF-α. Overall, the responses to the pool of CoV-2 S protein and individual peptides 163 and 164 were similar. T cell responses to 190/191 and 198/199 showed similar trends (**Supplemental Fig. S1**). Since the responding T cells are mostly CD4+, we conclude that peptides 163/164, 190/191, and 198/90 contain epitopes presented predominantly by MHC-II proteins.

**Figure 3:**
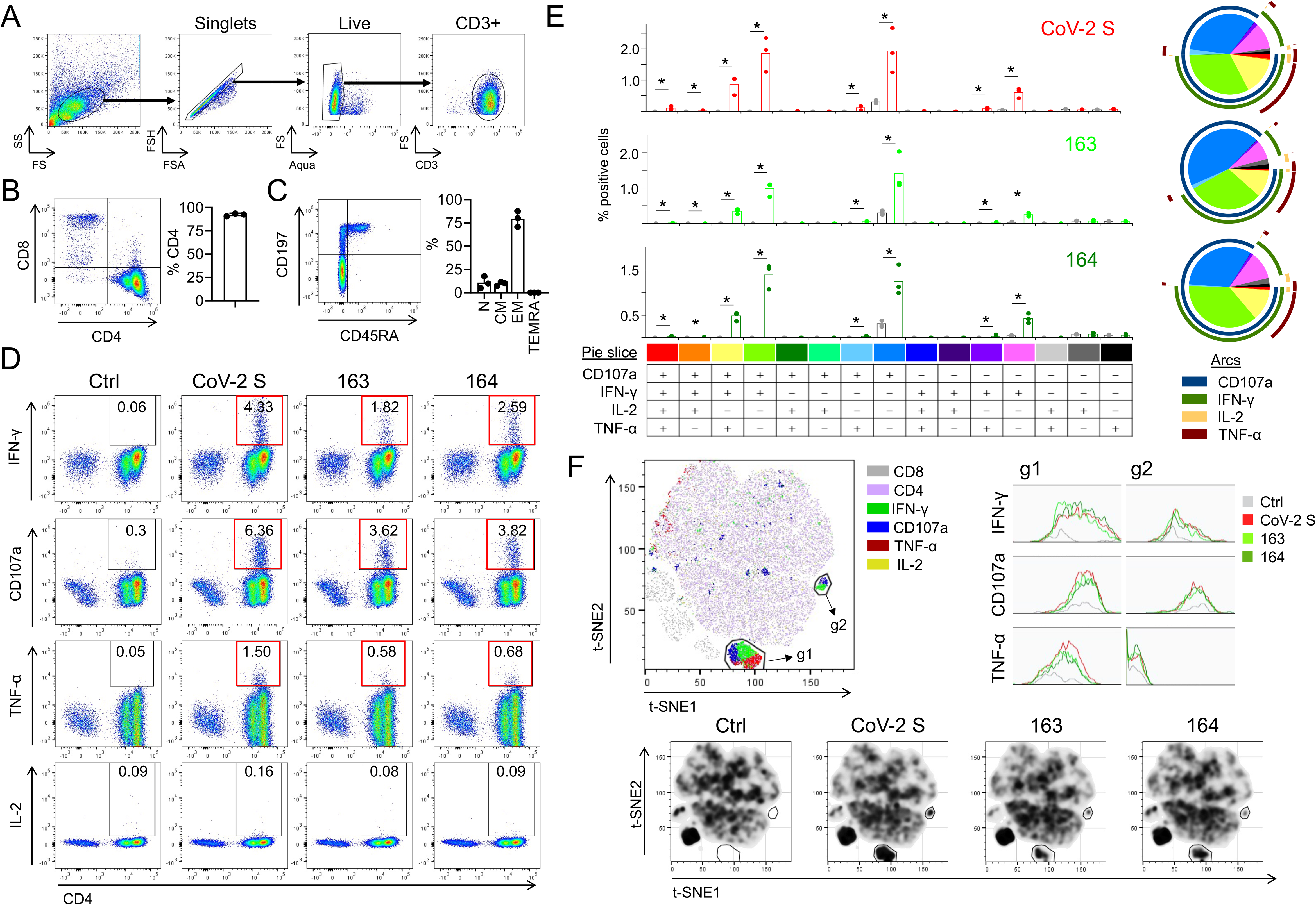
Functional characterization of in-vitro-expanded cross-reactive cells. A. Gating strategy. B. Representative dot plot for CD4/CD8 cells in the CD3+ population, and summary of CD4 expression in HCoV S pool in-vitro expanded cells from 3 donors. C. Representative dot plot for CD45RA/CD197 in the CD4+ population, and summary of the percentage of naïve (N), central memory (CM), effector-memory (EM), and EM re-expressing RA (TEMRA) populations in in-vitro expanded cells from 3 donors. D. Representative intracellular staining plots for IFN-γ, TNF-α, and IL-2 production, and CD107a mobilization to surface in the CD3+ population after re-stimulation of cross-reactive in-vitro expanded T cells with SARS-CoV-2 S pool (CoV-2 S) or individual peptides 163 and 164; positive responses shown in red boxes (>3-fold background). E. Visualization of the polyfunctional response using SPICE (Roederer et al., 2011): bar graph for each stimulating antigen (red for CoV-2, light green for 163 and dark green for 164) and comparison to control (grey) (p-values<0.05 using Wilcoxon Rank Sum Test are shown); pie and arcs graphs showing the combined contribution of each marker (pie slices’ colors correspond to colors shown at the bottom of bar graphs). F. t-SNE analysis of concatenated data from 3 donors for stimulation with CoV-2 S pool, peptides 163 and 164, and DMSO (Ctrl), showing density plots for each condition. Two gates (g1, g2) were drawn indicating major differences among stimulated and unstimulated samples. Histograms show the levels of IFN-γ, TNF-α, and CD107a in each gate for control (grey), CoV-2 S (red), 163 (light green) and 164 (dark green). Representative density plots for responses of d0801 are also shown.

### HLA restriction and epitope mapping

For epitope mapping studies we focused on the broadly-recognized overlapping peptide pair 163/164. Because of the possibility that multiple epitopes and/or multiple MHC molecules might be involved in the observed T cell responses, we generated a panel of T cell clones that recognize peptides 163/164, reasoning that HLA mapping and epitope identification would be facilitated by investigating T cell clones, which each would be expected to recognize a single epitope presented by a single MHC protein. Using in-vitro expanded cells from the same donors as investigated in Fig. 2 and screening for 163/164 reactivity, 49 clones were isolated and were tested for HLA restriction and defined epitope recognition (**Supplemental Table S2**). Donors d0801, d1102 and d1202 between them express fifteen different MHC-II alleles (**Supplemental Table 1**), of which ten alleles are predicted to bind at least one epitope within the 163/164 21-mer sequence (**Supplemental Table S3**). To begin to define HLA restriction for these clones, we used partially-HLA-matched antigen-presenting cells (APCs) and antibody-blocking experiments (Fig. 4A-D). Representative experiments using partially-HLA-matched APC are shown in Fig. 4A, and blocking experiments for the same clones using antibodies to HLA-DR, -DQ, -DP and HLA-A,B,C are shown in Fig. 4B. Summaries of such experiments for several clones are shown in Fig. 4C-4D, along with relevant HLA alleles for the donors and the partially-HLA-matched APC (Fig. 4E). For example, clone #143 from donor d0801 responded when peptides 163/164 were presented by LG2 cells, which share DQ5 (DQA1*01:01/DQB1*05:01) and DP4 (DPA1*01:03/DPB1*04:01) with the donor, but not when presented by 9068 cells, which share DR8 (DRB1*08:01), nor with single HLA-transfected DP4 cells (MN605), which all together suggests a DQ5 restriction (Fig. 4A left panels). This was confirmed by blocking antibody studies, where anti-DQ but not anti-DR, anti-DP, or anti-HLA-ABC was observed to inhibit the observed response (Fig. 4B, left panel). Similarly, clone #83 from donor d1202 showed strong reactivity to 9068 cells, which was blocked by anti-DP antibody; suggesting restriction by DP2 (DPA1*01:03 DPB1*02:01), the DP allele shared between the line and the donor (Fig. 4A-B, center panels). Clone #159 from donor d0801 showed reactivity to the DP4-transfected line and to a lesser extent to LG2 cells, which also express DP4; the blocking experiment confirmed the DP4 restriction (Fig. 4A-B, right panels). In this way, 13 of the clones derived from donor d0801 were categorized as DQ5-restricted and 2 were categorized as DP4-restricted, while 3 of the T cell clones derived from donor d1202 were categorized as being restricted by DP2 and one clone by DP4 (**Supplemental Table 2**). DP2 and DP4 are very similar proteins, with identical alpha subunits and only 4 amino acid changes in the beta subunit (V36A, D55A E56A E69K) (**Supplemental Fig. S2**), and two clones were observed to recognize 163/164 presented by either allele (d0801#120 and #159, **Supplemental Table S2**). We did not find clones restricted by DR alleles.

**Figure 4:**
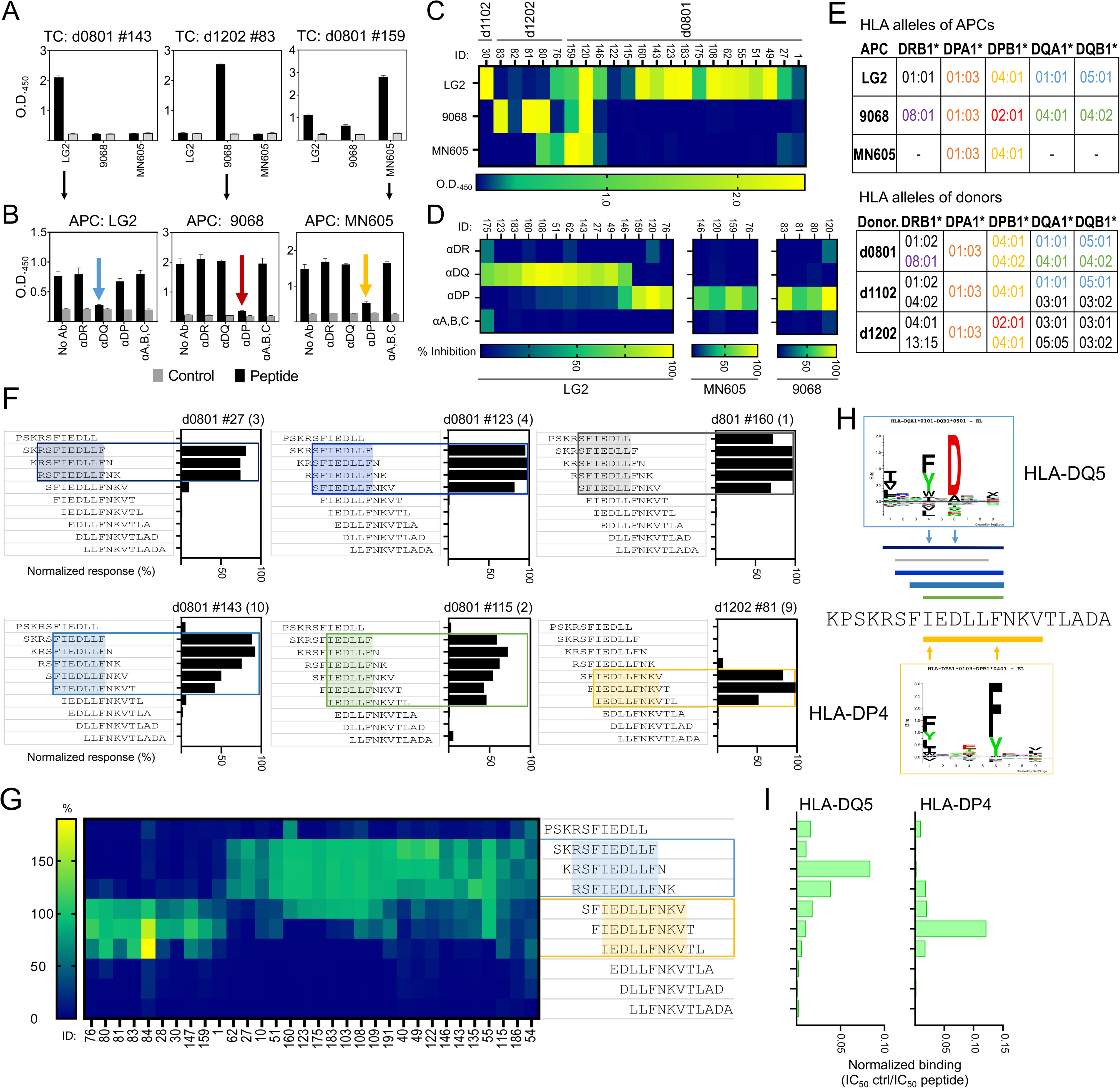
HLA restriction and epitope mapping. A. Responses of 3 representative T cell clones with confirmed reactivity to 163/164 presented by partially-match HLA cells (LG2, 9068, MN605), measured as IFN-γ in cell culture supernatant by ELISA; in black are responses to 163/164, and in gray are background responses (DMSO). B. Blocking of the responses in same 3 clones presented in A, using HLA-specific blocking antibodies to DR, DQ, DP and Class I; colored arrows show the inhibition. C. Summary of the responses to peptides 163/164 presented by the partially-match HLA cells in different clones (clone ID at top of graph); color scale represents the ΔOD_450_ (peptide minus DMSO). D. Summary of blocking experiments in different clones; color scale represents the percentage inhibition respect to no-antibody. E. Allele expressed by partially-match HLA cells that are shared with donors originating the T cell clones; shared alleles highlighted in colors. F. Responses to a set of truncated peptides (11-mers, overlapped by 10) covering the whole SARS-CoV-2 163/164 sequence, measured as IFN-γ in supernatant by ELISA; bars represent percentage of the response of each truncated peptide to the response of the full-length peptide. Representative clones for different reactivity patterns observed are shown; clone ID at top of graph with number of clones exhibiting a similar pattern in parenthesis. Minimal sequence required to explain reactivity is highlighted in each case. G. Summary of 32 clones analyzed; two partially overlapped main patterns can be observed, boxed in blue (for DQ) and yellow (for DP). H. Location of minimal epitopes from panel F shown aligned with the full-length 163/164 sequence (color of lines match color of box; thickness of line represents approximate frequency of the pattern). Predicted binding motifs for DQA1*01:01/DQB1*05:01 (top, DQ5) and DPA1*01:03/DPB1*04:01 (bottom, DP4) are shown as sequence logos. I. Normalized binding (IC_50_ control / IC_50_ peptide) of the set of truncated peptides to purified DQ5 and DP4 proteins.

To map the precise epitopes recognized by these clones, we evaluated the T cell response to a series of short peptides (11-mers overlapped by 10) covering the whole sequence present in the 163/164 peptides. We identified seven distinct patterns of the minimal peptide sequences that were required to activate the 29 clones analyzed (Fig. 4F; **Supplemental Table S2**), which segregated into two main groups sharing similar reactivity (Fig. 4G). One group, which consisted of clones that mapped to DQ5 (DQA1*01:01/DQB1*05:01) as defined by the presentation/blocking experiments, all responded to length variants of core epitope RSFIEDLLF (blue box in Fig. 4G). The other group, which consisted of clones that mapped to DP2 or DP4 (DPA1*01:03/DPB1*04:01 or DPA1*01:03/DPB1*02:01), all responded to length variants of a different core epitope, IEDLLFNKV (yellow box in Fig. 4G). DQ5-restricted clones recognized minimal peptide sequences 6-9 residues long, whereas the DP4-restricted clones recognized minimal peptide sequences 9 residues long (Fig. 4F, shaded regions, and Fig. 4H, lines above and below sequence). These recognition patterns are in good agreement with the binding predictions for these specific alleles (**Supplemental Table S3**), with minimal peptide sequences centered on the respective predicted core epitopes (Fig. 4H).

We validated tight binding of the 163/164 epitope to DQ5 and DP4, using purified proteins in fluorescent-peptide competition binding assays (**Supplemental Fig. S3**), and confirmed the differential core epitope selection by DQ5 and DP4 using the same set of minimal-length peptides as used to assess T cell recognition patterns (Fig. 4I). Maximal binding to DQ5 was observed for peptides with the core epitope RSFIEDLLF, whereas maximal binding to DP4 was observed for peptides with the core epitope IEDLLFNKV. Thus, DQ5 and DP4 both bind epitopes within the 163/164 sequence, but with a three-residue register shift between the core epitopes, consistent with the recognition patterns observed for the DQ5-, DP2-, and DP4-restricted T cell clones (Fig. 4H).

### Cross-reactive recognition of 163/164 variants from circulating HCoVs

The 163/164 epitope is highly conserved across the four circulating HCoVs, and we sought to determine the extent of T cell cross-reactivity between the corresponding epitopes in different viruses. To evaluate this, we tested responses of T cell clones from the COVID-19 donors to stimulation with overlapping peptides from the four HCoVs covering sequences corresponding to 163/164 in SARS-CoV-2 (Fig. 5A). All clones tested responded to a HCoV homolog from at least one virus (**Supplemental Table S2**). Responses by four representative T cell clones are shown in Fig. 5B. Hierarchical clustering of the responses of 17 clones to the different HCoV homolog peptides resulted in segregation of the clones into 4 major groups (Fig. 5C), which correspond to DQ5 or DP2/DP4 restriction patterns identified before. These groups show preferences for homologs from different viruses. For instance, clones in the purple group, categorized as DQ5-restricted, show reactivity to SARS-CoV-2, OC43, and HKU1, while clones in the turquoise group, also DQ5-restricted, show reactivity only to SARS-CoV-2 and HKU1. Clones in the gold and magenta groups, categorized as DP4 or DP2, show reactivity to SARS-CoV-2 and HKU1 (gold) or to SARS-CoV-2, NL63, and 229E (magenta).

**Figure 5:**
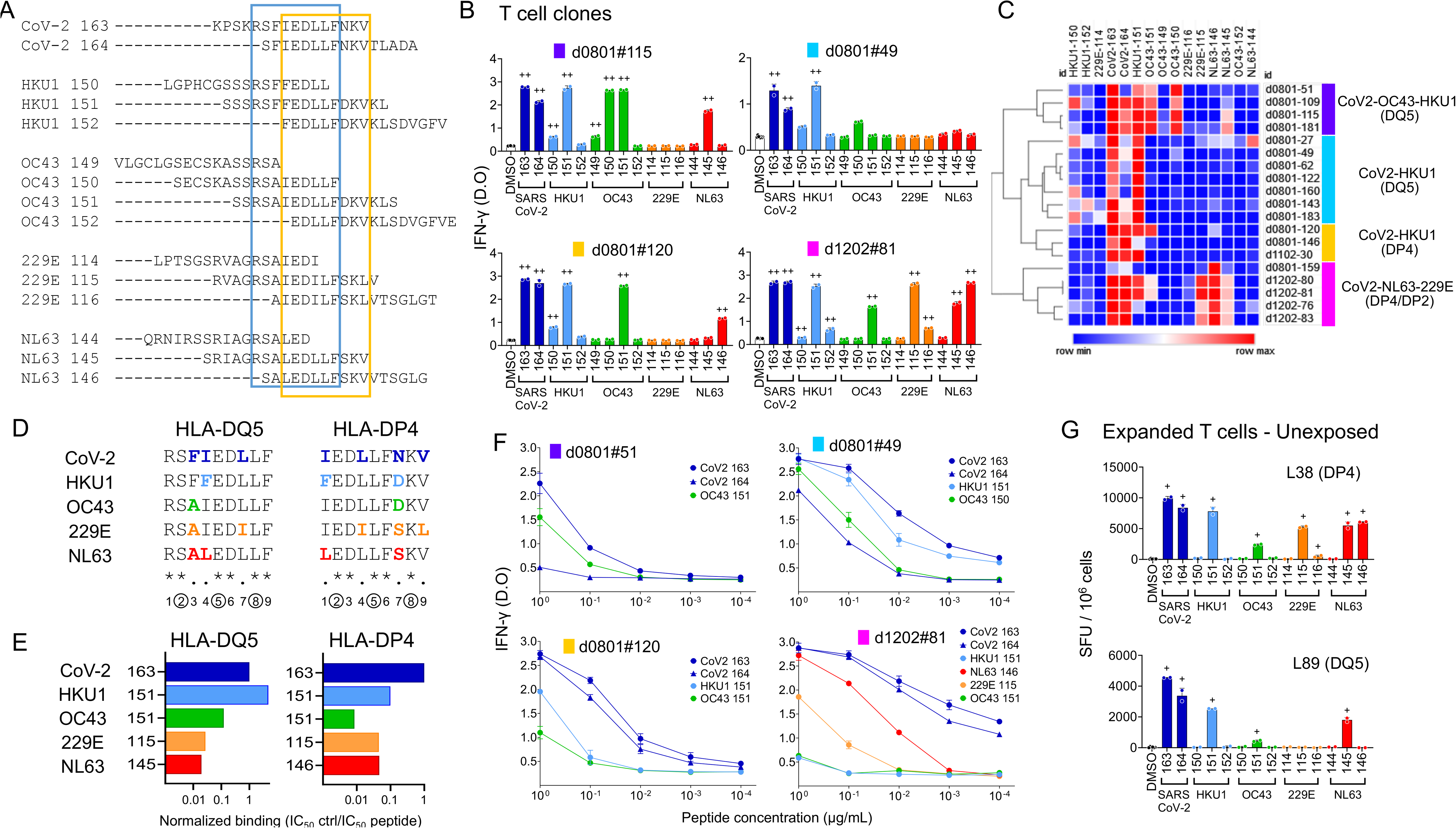
Cross-reactive recognition of 163/164 homologs from circulating HCoVs. A. Sequence of peptides 163/164 from SARS-CoV-2 and corresponding peptides in HKU1, OC43, 229E and NL63; boxes enclose the DQ5 (blue) and DP4 (yellow) core epitopes (see Fig. 4). B. Responses of representative T cell clones measured as IFN-γ in supernatant by ELISA. C. Hierarchical clustering of responses of 19 T cell clones to different homologs. Four major groups were defined: CoV2-OC43-HKU1/DQ5 (purple); CoV2-HKU1/ DQ5 (cyan); CoV2-HKU1/DP4 (gold); CoV2-NL63-229E/DP4-DP2 (magenta). D. Sequence alignment of DP4 and DQ5 epitope 9-mers of SARS-CoV-2 and homolog peptides from the four HCoVs. Numbers at the bottom indicate each position of the 9-mer and if it is identical (*) or there are changes (.) in the HCoVs relative to SARS-CoV-2 sequence; changes are highlighted in color in each sequence. E. Normalized binding (IC_50_ control / IC_50_ peptide) of homolog peptides from HCoVs at the peak of the response relative to binding of SARS-CoV-2 peptide 163, to purified HLA-DQ5 and DP4 proteins in an in-vitro competition assay. F. Dose-response of selected T cell clones to homolog peptides, measured as IFN-γ in supernatant by ELISA. G. Responses of in-vitro expanded cells from unexposed donors (expanded with SARS-CoV-2 peptides) to homolog peptides, measured as IFN-γ ELISpot (+ indicates positive response by DFR2X); indicated in parenthesis is the relevant HLA present in each donor.

These response patterns can be understood by considering the 163/164 epitope sequences in the various viruses. Alignment of the SARS-CoV-2 and HCoV sequences in the region of the DQ5 and DP4 core epitopes shows that for both binding registers, the P2, P5, and P8 positions are 100% conserved (Fig. 5D). These positions are located where the major T cell contacts are expected for conventionally oriented T cell receptors (Rossjohn et al., 2015; Stern and Wiley, 1994). Conservation of the key TCR contact residues helps to explain the overall high degree of SARS-CoV-2 – HCoV cross-reactivity in these epitopes. Residues at other positions are less conserved in both DQ5 and DP4 registers, contributing to the differential MHC binding that we observed in the in vitro competition binding assay (Fig. 5E). DQ5 showed a preference for the HKU1 homolog, which it binds more strongly than the SARS-CoV-2 homolog, with reduced binding to the OC43 homolog and substantially weaker binding to the NL63 and 229E homologs. For DP4, the strongest preference among the HCoV peptides again was for the HKU1 homolog, followed by 229E and NL63, and very little binding for OC43. These patterns could be understood in terms of substitutions at the positions expected to bind into the major MHC side-chain binding pockets at P1, P4, P6, or P9, and the minor pockets or “shelves” at P3 and P7. For example the improved DQ5 binding of the HKU1 homolog relative to SARS-CoV-2 apparently is due to Phe at the P4 position, which is the only difference between the SARS-CoV-2 and HKU1 core epitope sequences, and a preferred residue at the P4 position (Fig. 4H). Similarly, the reduced binding of OC43, 229E, and NL63 homologs appears to be due to combinations of effects at the P3, P4, and P7 positions. For DP4, weaker binding of the OC43 homolog appears to be due to an Asn-to-Asp substitution at the P7 “shelf” position, which is partially compensated in the HKU1 homolog by Phe at the preferred DP4 P1 anchor residue position (Sidney et al., 2010b). These binding differences also help to explain the reactivity groups defined in Fig. 5C. The purple and cyan groups, categorized as DQ5, show little or no reactivity with 229E and NL63, which exhibited minimal DQ5 binding for their 163/164 homologs. Similarly the gold and magenta groups, categorized as DP4, exclude OC43, which was worst DP4 binder. The restricted specificity for SARS-CoV-2 and HKU1 in the cyan group probably indicates an important TCR preference for Phe over Ala at the “shelf” P3 position in the DQ5 register (Sidney et al., 2010a), and in the gold group an important TCR contact at the Asn/Asp at P7 in the DP4 register.

Dose-response experiments show that the preferred homolog reactivity is, in most cases, comparable to the reactivity to the SARS-CoV-2 peptide over a wide range of concentrations (Fig. 5F). For instance, for clone #51 in the purple group (DQ5, CoV-2/OC43/HKU1), dose-dependent reactivities to SARS-CoV-2-163 and OC43-151 peptides are similar. Likewise, for clone #49 in the cyan group (DQ5, CoV-2/HKU1) dose-dependent reactivities for SARS-CoV-2-163 and HKU1-151 also are similar. For clone #120 in the gold group (DP4, CoV-2/HKU1) and clone #81 in the magenta group (DP4/DP2, CoV-2/NL63/229E), reactivities for the targeted HKU1 and NL63 peptides were approximately 10-fold weaker than for SARS-CoV-2.

Finally, we performed similar experiments using cross-reactive T cells from unexposed donors obtained by expansion with SARS-CoV-2 163/164 (Fig. 5G). The patterns of cross-reactivity observed for these donors were similar to those defined for the clones derived from COVID-19 donors, with donor L38 exhibiting a pattern similar to the magenta group, and L89 similar to the gold group.

### Broad recognition of 163/164 in the population

We evaluated the frequency of response to 163/164 in additional unexposed donors, in order to assess the presence of a cross-reactive response before COVID-19, and in donors receiving mRNA-based COVID-19 vaccines, to assess if vaccination can induce responses to this epitope. For the unexposed donors we used PBMCs cryopreserved between 2015-2018. As before we measured IFN-γ T cell responses directly ex-vivo and after expansion with HCoV S pools or SARS-CoV-2 163/164 peptides. Out of 9 donors analyzed, we found positive responses detectable in direct ex vivo assay in 2 donors; in short-term T cell cultures expanded with HCoV S pools we found positive responses in 4 donors, and in cultures expanded with SARS-CoV-2 163/164 peptides, we found positive responses in 8 donors (Fig. 6A). We also measured responses in individuals who had received one of the available mRNA vaccines. Of 15 vaccinated donors tested we observed responses in direct ex vivo assays in 8 donors, including those with no evidence of previous COVID-19 infection (2 positive of 5 tested), individuals that had COVID-19 and were later vaccinated (3 postive of 5 tested), and individuals that were vaccinated and later got COVID-19, i.e. breakthrough cases (3 positive of 4 tested) (Fig. 6B). As noted above in Fig 2D, for convalescent COVID-19 donors 7 out of 11 exhibited positive IFN-γ T cell responses to 163/164 in direct ex vivo assays. Thus, the SARS-CoV-2 163/164 epitope is recognized broadly in unexposed, mRNA-vaccinated, and COVID-19 donors.

**Figure 6:**
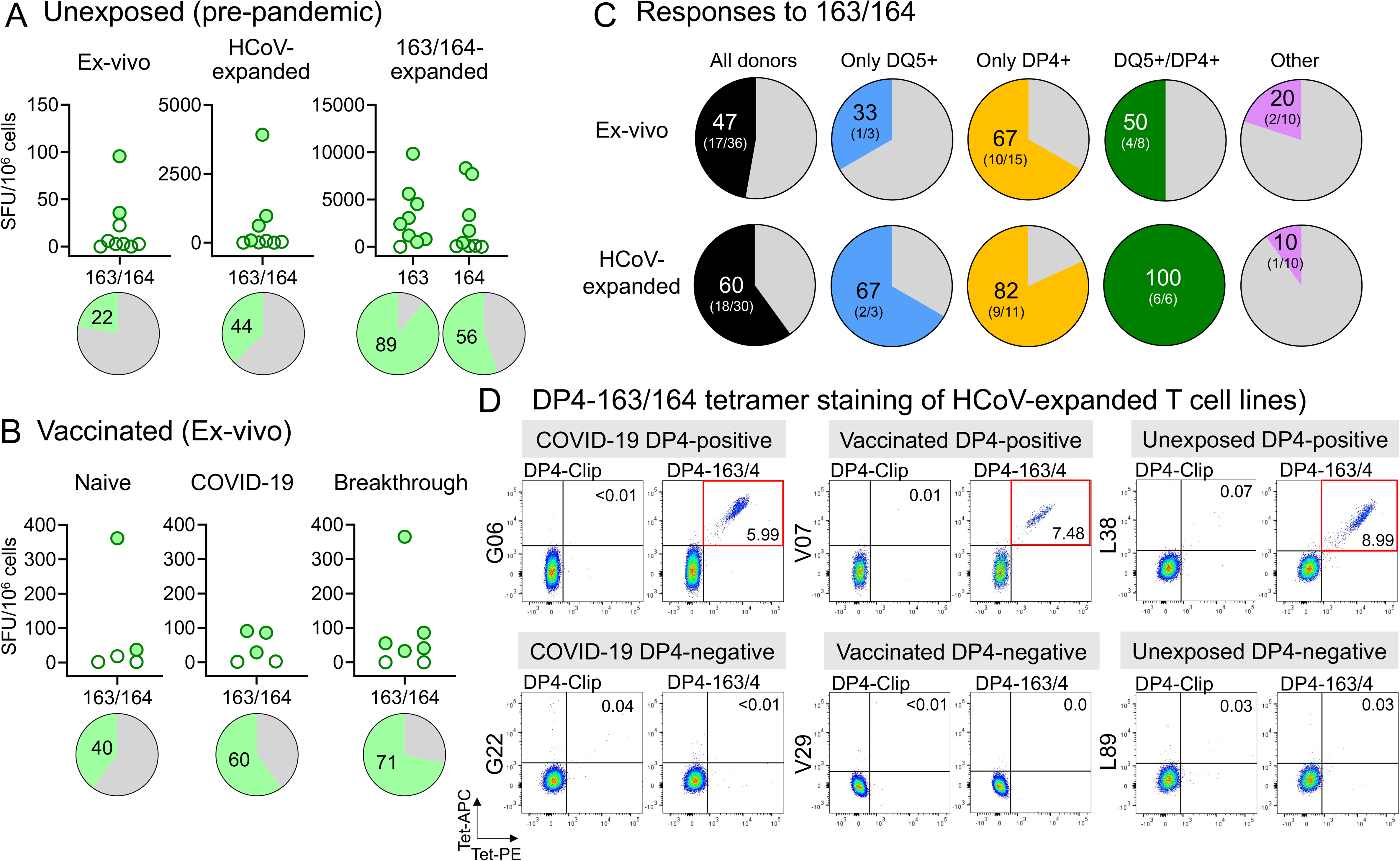
Broad recognition of 163/164 in the population. Responses to peptides 163/164 in 9 pre-pandemic donors, ex-vivo and after in-vitro expansion with HCoV S pools (HCoV-expanded) or SARS-CoV-2 peptides 163/164 (163/164-expanded). B. Ex-vivo responses to peptides 163/164 in vaccinated donors (naïve: vaccine recipients, without previous COVID-19; COVID-19: vaccine recipients, with previous COVID-19; breakthrough: vaccine recipient with COVID-19 after vaccination). In A and B percentage of positive responses are shown in the pie graphs. C. Responses to 163/164 ex-vivo and in in-vitro expanded cells (HCoV-expanded) for donors categorized according to HLA-DQ5 and HLA-DP4 status: “All donors” groups donors regardless DQ5/DP4 status; only DQ5 express DQ5 but not DP4; only DP4 express DP4 but not DQ5; DQ5/DP4 express both DQ5 and DP4; other express neither DQ5 nor DP4. Percentage of donors with a positive response in each group is shown (number of positive and total donors in parenthesis). D. DPA1*01:03/DPB1*04:01-163/164 tetramer staining of in-vitro HCoV S pool expanded T cells from COVID-19, unexposed, and vaccine recipients; representative staining in one donor expressing DP4 (top) and one donor not expressing DP4 (bottom) in each group. Double-tetramer (PE and APC) staining in CD4+ population is shown in dot plots; DP4-Clip tetramers used as controls.

We analyzed the T cell responses to 163/164 from all of the donors, considering their DQ5 and DP4 status (Fig. 6C). For those who expressed only DQ5 but not DP4, 33% responded to peptide 163/164 in direct ex vivo assays, and 67% after in vitro expansion. For those who expressed only DP4 but not DQ5, the percentage responding was substantially larger, 67% in direct ex vivo assays, and 82% after expansion. In this analysis, we included also 3 donors with the closely-related DP4 variant DPA1*01:03/DPB1*04:02 (DP402), and 6 donors expressing both DPB1*04:01 and *04:02 variants. The three amino acid differences between DP4 and DP402 are buried underneath the peptide largely away from peptide binding pockets (**Supplemental Fig. S2**), and known to have little if any effect on peptide binding specificity (Castelli et al., 2002). Overall, this confirms that 163/164 can be considered as a broadly-recognized immunodominant epitope in individuals expressing DQ5 or DP4 alleles.

### Tetramer staining

We investigated the use of MHC-II tetramers to following cross-reactive T cell populations recognizing the 163/164 eptiope. We focused on DP4 because of the high prevalence of this allele across most human population groups ((Castelli et al., 2002; Sidney et al., 2010b), **Supplemental Fig. S2**). We used tetramers carrying a 15-mer peptide centered around the DP4-restricted 163/164 epitope RSF<u>IEDLLFNKV</u>TLA (core epitope underlined), labeled with APC or PE fluorophors. We first tested for specific recognition using T cell clones recognizing the 163/164 epitope presented by DP4 (clone d801 #120, 146, 159 and d1202 #76), DP2 (clone d1202 #81 and 83) and DQ5 (clone d801 #143, 51, 62 and 49) (**Supplemental Fig. S4A**). The DP4-restricted T cell clones were recognized as a distinct population strongly staining with both PE- and APC-labeled tetramers, while the other clones, as well as a negative control DP4-Clip tetramer, exhibited no detectable staining. T-cell lines from DP4-positive donors expanded in vitro using HCoV S pools exhibited populations staining strongly with DP4-163/164 tetramer as compared to negative control, including lines from a convalescent COVID-19 donor (G06), a vaccinated donor (V07), and an unexposed donor (L38) (Fig. 6D). For the same unexposed donor, we evaluated DP4-163/164 tetramer staining in resting unstimulated PBMC, and observed a small population visible at ~5-fold increased abundance over the non-specific staining background (**Supplemental Fig. S4B**). Expanded T cell lines from unexposed and vaccinated DP402 donors also could be detected (**Supplemental Fig. S4C**). Finally, we evaluated DP4-163/164 tetramer staining for T cell populations expanded in vitro using SARS-CoV-2 163/164 or the individual HCoV homologs of that epitope (**Supplemental Fig. S5A**). DP4-163/164 tetramer-positive populations were observed for T cell lines expanded with each of the homologs, with the relative size of the populations matching ELISpot results on these same lines (**Supplemental Fig. S5B**), and consistent with the patterns of HCoV reactivity observed in Fig. 5. These results confirm DP4 presentation of the 163/164 peptide as identified in cellular and biochemical studies, and validate the use of the DP4-163/164 tetramer in detecting SARS-CoV-2 and HCoV-cross-reactive T cell populations.

### T cell receptors

To further characterize the cross-reactive response, we analyzed the TCR repertoires of T cell populations responding to homologous SARS-CoV-2 and HCoV antigens. To identify TCRα and TCRβ sequences of DP4-specific T cells recognizing the 163/164 epitope, we sorted DP4-163/164 tetramer-positive cells from two COVID-19 and two pre-pandemic donors, all DP4-positive, after in vitro expansion with SARS-CoV-2 163/164. A total of 173 TCRα and 184 TCRβ unique sequences were identified (**Supplemental Table S5**). Limited TRAV and TRBV gene sharing among the donors was observed (Fig. 7A), with diverse CDR3 sequences (Fig. 7B). To extend these results to additional cross-reactive specificites beyond 163/164, we analyzed bulk TCRα and TCRβ repertoires from 6 COVID-19 donors after expansion with HCoV S pools. A total of 1,663 TCRα and 2,177 TCRβ unique sequences were identified in these 6 samples (**Supplemental Table S6**). We also sequenced TCRα and TCRβ repertoires from one pre-pandemic donor after expansion with HCoV S pools (**Supplemental Table S6**). Many of the CDR3 sequences identified in these polyclonal lines were also observed in DP4-163/164-tetramer sorted cells or T cell clones derived from the same donors. Several TCRα and TCRβ clonotypes were shared among two or three donors, including some also observed in DP4-163/164-tetramer sorted populations (Fig. 7C, **Supplemental Table S7**), and in previously reported datasets of COVID-19-associated TCRβ reperotires (**Supplemental Table S8**; (Dykema et al., 2021; Low et al., 2021; Nolan et al., 2020). We used the GLIPH algorithm (Glanville et al., 2017) to cluster TCRβs by shared specificity and identify sequence motifs that might be shared more broadly between donors than exact CDR3 sequence matches. Analysis of the pooled TCRβ sequences from expanded polyclonal lines, T cell clones, and DP4-sorted cells revealed 117 TCR convergence groups, some of which were observed across multiple donors (**Supplemental Table S9**). Among them, two clusters that could be associated to DP4 were observed (Fig. 7D). Each was observed in 6 different donors, including samples from COVID-19 and pre-pandemic polyclonal lines and DP4-163/164-tetramer sorted cells. One cluster that could be associated to DQ5 was also observed in samples from 3 donors as well as DQ5-restricted clones (Fig. 7D). Some of these clusters also were detected in previously reported TCRβ repertoires (Nolan et al., 2020), (Low et al., 2021), and (Dykema et al., 2021). Finally, we defined candidate TCRα and TCRβ pairings, using sequence information from selected T cell clones derived from two COVID-19 donors responsive to peptides 163/164. Four candidate TCRα/TCRβ pairs for DQ5 and one for DP4 were assigned by detection in multiple clones, with four candidate TCRα/TCRβ pairs for DP2 and an additional one for DP4 also assigned but not confirmed in multiple clones (Fig. 7E). In summary, we find a highly diverse repertoire of TCRs recognizing the 163/164 peptide in the context of DP4 and other HLA alleles, with little sharing of either TRAV/TRBV gene usage or CDR3 sequences between donors, although many low-frequency public TCRα and TCRβ clonotypes and convergence groups could be indentifed.

**Figure 7:**
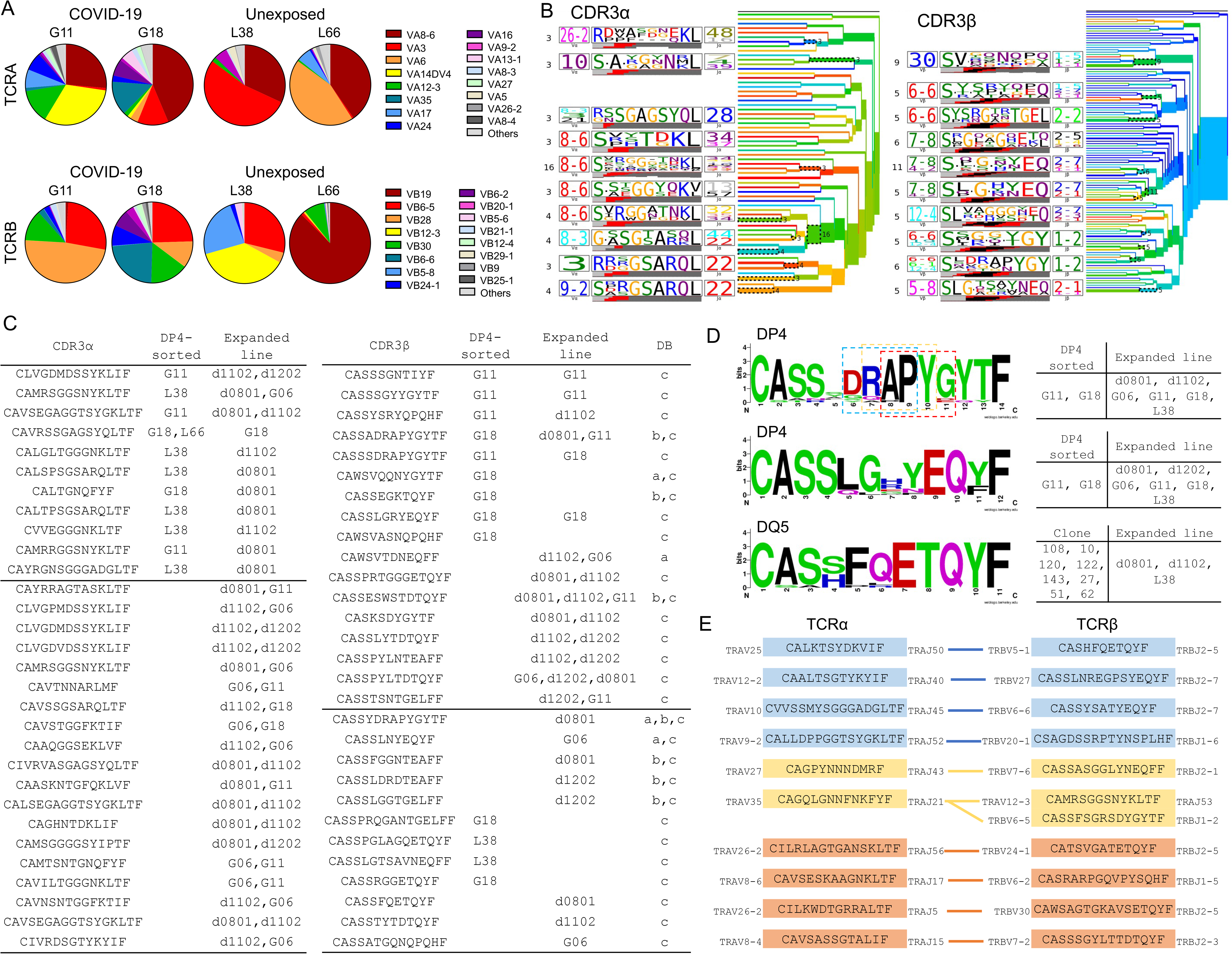
TCR repertoires: A. TCR Vα and Vβ usage in DP4-163/164 tetramer-sorted cells expanded from DP4+ COVID-19 and unexposed donors with peptides 163/164. B. Clustering tree of CDR3α and CDR3β clonotypes and sequence logo in the sorted cells from the four donors combined, obtained using TCRDist algorithm (Dash et al., 2017). C. Summary of public CDR3α and CDR3β clonotypes identified in DP4-sorted samples, in-vitro HCoV S pool expanded lines (unsorted), and T cell clones; sequences identified in reported datasets are indicated: (a) Nolan et al., (Nolan et al., 2020), ( b) Low et al., (Low et al., 2021), (c) Dykema et al. (Dykema et al., 2021). D. Sequence logos of three selected GLIPH clusters (Glanville et al., 2017) for CDR3β clonotypes from DP4-163/164-sorted, HCoV S pool-expanded T cells, and T cell clones. CRG final scores: 2.8×10^−14^ (top), 4.4×10^−14^ (middle), and 2.5×10^−15^ (bottom); significant motifs in top cluster indicated in boxes (RAPY (13 sequences, p<0.001), APYG (15 sequences, p<0.001), DRAP (13 sequences, p<0.001)). E. Candidate DQ5 (blue), DP4 (yellow), and DP2 (orange) TCRα/β pairs inferred from sequencing of T cell clones.

## Discussion

We studied T cell cross-reactivity between SARS-CoV-2 spike protein and the four circulating seasonal coronaviruses by measuring the response to SARS-CoV-2 and homolgous HCoV spike peptides in peripheral blood samples from convalescent COVID-19 donors, vaccine recipients, and individuals not exposed to SARS-CoV-2. We challenged T cells from SARS-CoV-2-exposed donors with HCoV spike peptide pools, and vice versa, making particular use of T cell lines from SARS-CoV-2 donors expanded in vitro by stimulation with HCoV spike peptides, which enriches for cross-reactive T cells. We identified several peptides recognized by cross-reactive T cell populations, in particular peptide 163/164, which dominated the spike cross-reactive response, and was recognized by T cell responses in most donors tested. The 163/164 region, located proximal to the S2’ cleavage site at the start of the fusion peptide, is highly conserved across SARS-CoV-2 variants and human coronaviruses, because processing at the spike S2/S2’ junction is necessary to release the fusion peptide essential for viral entry and membrane fusion. We used a panel of T cell clones to identify minimal epitopes and presenting MHC molecules, and identified two major patterns of reactivity, with an N-terminal epitope RSFIEDLLF S_815-823_ presented by DQ5, and a partially-overlaping C-terminal epitope IEDLLFNKV S_818-826_ presented by DP2 and DP4. T cells recognizing these eptiopes were highly cross-reactive for corresponding SARS-CoV-2 and HCoV sequences.

Most donors tested recognized the 163/164 peptide, including severe and mild COVID-19 donors, individuals receiving mRNA COVID-19 vaccines, and previously unexposed donors, although most previously unexposed donors required in vitro expansion to increase responding T cell populations to detectable levels. The broad recognition of these epitopes is driven by the prevalence of the presenting MHC molecules, particularly HLA-DP4, which is the most common MHC allele worldwide (Castelli et al., 2002; Sidney et al., 2010b). The same epitope was presented by DP4 and also by the closely related alleles DP2 and DP402, which share similar peptide binding motifs, identical DPα subunits and DPβ subunits each with only four substitutions (**Supplemental Fig. S2**). Between them these alleles cover a large fraction of many human populations worldwide (**Supplemental Fig. S2**).

The key to this cross-reactive recognition seems to be the remarkable conservation across human coronaviruses of identical amino acids at expected T cell contact positions for both the DQ5 and DP4 binding frames. The selection for DP4 and DQ5 as preferred presenting elements for cross-reactive recognition of SARS-CoV-2 and HCoV homologs may be related to their particular binding motifs, which accommodate peptide sequence variability while still presenting identical residues for TCR recognition. Other MHC-II proteins that present overlapping peptides from this region in different binding frames would not present such a conserved set of residues. For example, a SARS-CoV-2 peptide overlapping 163/164 was identified in a CD4 T cell epitope screen (Verhagen et al., 2021) and as a naturally-processed epitope derived from SARS-CoV-2 spike protein (Parker et al., 2021), where it was predicted to be presented by DR3 and DR4, respectively, but for these alleles the expected T cell contact positions are not conserved.

We observed that T cell populations cross-reactive for both SARS-CoV-2 and seasonal HCoVs comprise only a small fraction of the overall response in both unexposed and COVID-19 donors. These responses were characterized by a highly diverse cross-reactive TCR repertoire, mainly specific to individual donors, although a few shared or public clonotypes were present across donors with varying abundance. There does seem to be a large repertoire of cross-reactive cells available for expansion, with a highly diverse CDR3 repertoire. Why cross-reactive T cells present before SARS-CoV-2 infection do not expand and dominate the overall response is not clear.

Immune reponses to the 163/164 region of the SARS-CoV-2 spike protein have been observed in other studies (Deng et al., 2021; Dykema et al., 2021; Low et al., 2021; Loyal et al., 2021; Mateus et al., 2020; Saini et al., 2021; Tarke et al., 2021; Woldemeskel et al., 2021). An overlapping epitope accessible in the pre-fusion conformation was recognized by antibodies from COVID-19 donors as one of the most highly recognized linear epitopes, and antibodies recognizing this sequence were detected in both COVID-19 and unexposed donors (Poh et al., 2020; Shrock et al., 2020; Voss et al., 2021). Several studies of CD4+ and CD8+ T cell responses in SARS-CoV-2 donors, including unbiased epitope screens as well as those based on MHC-binding predictions, identified peptides overlapping the S_811-831_ sequence among many others (Deng et al., 2021; Dykema et al., 2021; Low et al., 2021; Saini et al., 2021; Tarke et al., 2021; Woldemeskel et al., 2021). A peptide overlapping the S_811-831_ sequence was found among MHC-II-bound peptides eluted from human monocyte-derived DCs pulsed in vitro with spike protein (Knierman et al., 2020), showing that this epitope is presented after antigen processing and presentation in a natural context, although in that study the presenting HLA molecules were not assigned (Knierman et al., 2020). The 163/164 region also has been investigated previously in the context of cross-reactivity between SARS-CoV-2 and HCoVs. Mateus et al indentified an overlapping epitope in one of 9 peptides for which cross-reactive T cell responses were validated for SARS-CoV-2 and HCoV variants (Mateus et al., 2020). Loyal et al. observed T cell populations responding to two overlapping peptides from the same region that we report here, in a study of SARS-CoV-2 epitopes recognized in uninfected donors, where they were shown also to contribute in the initial response to primary SARS-CoV-2 infection (Loyal et al., 2021). An epitope overlapping with 163/164 was one of two highlighted in a recently study by Low et al.,who mapped the specificity, HLA restriction, and HCoV cross-reactivitiy of a set of 247 T cell clones isolated from 22 COVID-19 and unexposed donors (Low et al., 2021). Dykema et al., mapped peptide reactivity for TCRs over-represented in cross-reactive, in-vitro expanded, CD4 T cell lines, and identified a sequence from the 163/164 region recognized by five TCR transfectants that also recognized the NL63 homolog (Dykema et al., 2021). In contrast to these studies, we systematically evaluated which epitopes dominated the SARS-CoV-2 – HCoV cross-reactive T cell response, we validated a proposed DP4 restriction through direct MHC-peptide binding and minimal peptide mapping studies, we identified DQ5 as a new important presenting molecule that recognizes a register-shifted epitope, and we identified patterns of cross-reactivity with the various HCoV homologs.

There are several limitations of our study. We mapped the specificity of the cross-reactive response by following IFN-γ-secreting cells, but non-IFN-γ-secreting populations could also contribute to the response. In expanded T cell lines we observed higher frequencies of T cells staining with DP4-163/164 tetramer than responding to the same peptide in IFN-γ ELISPot essays, indicating that some T cells can recognize the epitope but not secrete IFN-γ. We observed the cross-reactive T cell response to involve mostly CD4^+^ T cells. This might be due to in vitro culture conditions that favor CD4^+^ over CD8^+^ T cell populations, or an intrinsic bias of cross-reactive T cells because of the different patterns of pMHC-TCR interaction for MHC-I and MHC-II proteins. We studied a relatively small group of 27 individuals exposed to SARS-CoV-2 antigens by infection or vaccination, mostly over 40 years of age. Younger individuals with more frequent previous exposures to HCoVs might show a different pattern of response. Our initial screen for immunodominant epitopes involved only three donors, all of whom recognized 163/164, but other immudominant cross-reactive epitopes might have escaped our attention, including those recognized by other MHC proteins. For all of the donors, previous HCoV infection was inferred but not observed, and we did not attempt to determine which donors were exposed previously to which of the HCoVs.

In conclusion, we identified a pan-coronavirus epitope that dominates the cross-reactive T cell response to the spike protein after SARS-CoV-2 infection. The epitope is highly conserved across human coronoaviruses, with T cell receptor contact positions invariant in each of two partially overlapping MHC-II binding frames. Most people will have CD4^+^ T cells responding to this epitope before SARS-CoV-2 infection, because of its robust presentation by common HLA molecules and the seasonal prevalence of infection by HCoVs. Responding T cells appear to be functionally competent and are strongly expanded ex vivo by cross-stimulation, but do not dominate the primary response after natural infection, at least as assessed 3-9 months post-infection. The S_811-831_ sequence, completely conserved in SARS-CoV-2 variants including delta and omicron, may be useful in studies relating pre-existing HCoV immunity to COVID-19 severity or incidence, and might be considered for inclusion in pan-coronavirus vaccination strategies.

## Acknowledgements

We acknowledge the contributions of people who donated blood samples, without whom this study would not have been possible. Robert W. Finberg passed away during the course of this work. We thank the UMass Memorial Medical Center Histocompoatiblity Laboratory and Jeff Bailey (Brown University) for assistance with HLA typing. We thank Mollie Jurewicz for assistance with TCR repertoire methodology, Manuel Garber for assistance with TCR repertoire analysis, and Ann M Rothstein for helpful discussions. We acknowledge the assistance of the UMass Chan Medical School Clinical Research Center, Flow Cytometry Core, and Deep Sequencing Core, and we thank the NIH Tetramer Core Facility (contract number 75N93020D00005) for providing DP4-163/164 tetramers. This work was supported by grants from the Massachusetts Consortium Pathogen Readiness MassCPR (AMM) and the UMass COVID-19 Pandemic Fund (LJS).

## Author Contributions

JMCC, ABA, and LJS concieved the analysis. RWF, MC, and AMM recruited donors and participated in study design. JC, CF, JMCC, and ABA assisisted with blood processing. ABA and JMCC performed all T cell studies. CF and AMM measured antibody responses. LL and GCW prepared recombinant proteins, and PNN performed MHC-peptide binding assays. ABA, JMCC, and LJS analyzed T cell data, HCoV homologies, peptide binding patterns, and TCR repertories. ABA, JMCC, and LJS wrote the manuscript.

## STAR Methods

### RESOURCE AVAILABILITY

#### Lead contact

Further information and requests for resources and reagents should be directed to lead contact.

#### Materials availability

This study did not generate new unique reagents.

#### Data and code availability

Any additional information required to reanalyze the data reported in this paper is available from the lead contact upon request.

### EXPERIMENTAL MODEL AND SUBJECT DETAILS

#### Blood, PBMCs and HLA typing

Whole blood from COVID-19 convalescent donors, healthy donors, or vaccine recipients was collected under protocol approved by the UMass Chan Medical School Institutional Review Board of the University of Massachusetts and informed consent was obtained from all subjects. Leukopaks were obtained from New York Biologics, Inc. (Southampton, NY). Peripheral blood mononuclear cells (PBMCs) were isolated using Ficoll-Paque (Cytiva, Marlborough, MA) density gradient centrifugation and used fresh or frozen until use. The HLA class II haplotype of pre-pandemic donors was determined using the Protrans HLA typing kits (Protrans Medizinische Diagnostische Produkte GmbH, Hockenheim, Germany) or The Sequencing Center (Fort Collins, CO); for other donors, HLA typing was performed using a Nanopore protocol (Stockton et al., 2020) or by the Histocompatibility Laboratory at UMass Memorial Medical Center (Worcester, MA).

### METHOD DETAILS

#### Generation of peptide-expanded T cells

Peptide-pool or individual peptide expanded T cell lines were generated for each donor by a single in-vitro expansion of freshly isolated or frozen PBMCs (2 x10^6^ cells in 1 mL in a 24 well plate) with a final concentration of 1 μg/mL of peptide. As antigens were used individual peptides; peptides covering the entire SARS-CoV-2 spike protein in a single pool or pools of 10 peptides; peptides pools covering the entire spike proteins of OC43, HKU1, NL63, and 229E. Cells were maintained in complete RPMI (CRPMI, RPMI 1640 supplemented with 10% AB+ human serum (GeminiBio, West Sacramento, CA), 50 μM beta-mercaptoethanol, 1 mM non-essential amino acids, 1 mM sodium pyruvate and 100 U/mL penicillin and 100 mg/mL streptomycin (all Gibco, Grand Island, NY)). After 3 days, cultures were supplemented with recombinant human IL-2 (Proleukine, Prometheus, San Diego, CA) at a final concentration of 100 U/mL. During the following 2-15 days, one-half of the medium was replaced with fresh CRPMI supplemented with 100 U/mL IL-2 every 3 days. When cultures reached confluence, cells were resuspended and one-half of the culture transfer to another well and fresh CRPMI+100 U/mL IL-2 added to replenish the original volume.

#### ELISpot assay

IFN-γ ELISpot were performed using Human IFN gamma ELISpot KIT (Invitrogen, San Diego, CA) and MultiScreen Immobilon-P 96 well filtration plates (EMD Millipore, Burlington, MA), following manufacturer’s instructions. Assays were performed in CST™ OpTmizer™ T cell medium (Gibco, Grand Island, NY). Peptides or peptides pools were used at a final concentration of 1 μg/mL per peptide; as negative controls were used DMSO (DMSO, Fisher Scientific, Hampton, NH) and a pool of human self-peptides (Self-1, (Becerra-Artiles et al., 2019)), and PHA-M (Gibco, Grand Island, NY) was used as positive control. For exvivo assays, PBMCs (~2-5×10^5^ per well) were incubated with peptides or controls for ~24-48 hours. For assays with cells expanded in-vitro, 2-5×10^4^ cells per well were incubated with an equal number of autologous irradiated PBMCs in the presence of peptides or controls for ~18 hours. Two to four wells of each peptide, pool of peptides, or PHA-M, and at least 6 wells for DMSO were usually tested. Secreted IFN-γ was detected following manufacturer’s protocol. Plates were analyzed using the CTL ImmunoSpot Image Analyzer (ImmunoSpot, Cleveland, OH) and ImmunoSpot 7 software.

#### Intracellular cytokine staining (ICS)

ICS was performed using in-vitro expanded T cells as previously described (Becerra-Artiles et al., 2019) with minor modifications. Briefly, autologous irradiated PBMCs were resuspended in CRPMI (w/o phenol red) +10% fetal bovine serum (FBS, R&D Systems) containing 1 μg/mL of each peptide and incubated overnight. The day of the assay, T cell lines were collected, washed and resuspended in the same medium and added to the pulsed PBMCs (1:1 ratio, usually 0.3×10^6^ cells each); at this time, anti-CD107a-CF594 (H4A3) was added, along with brefelding A and monesin at the suggested concentrations (Golgi plug / Golgi stop, BD Biosciences, San Jose, CA). After 6 hours incubation, cells were collected, washed, and stained using a standard protocol, which included: staining for dead cells with Live/Dead Fixable Aqua Dead Cell Stain Kit™ (Life Technologies, Thermo Fisher Scientific, Waltham, MA); blocking of Fc receptors with human Ig (Sigma-Aldrich, St. Louis, MO); surface staining with mouse anti-human CD3-APC-H7 (SK7), CD4-PerCPCy5.5 (RPA-T4), CD8-APC-R700 (RPA-T8), CD14-BV510 (MϕP9), CD19-BV510 (SJ25C1), CD56-BV510 (NCAM16.2); fixation and permeabilization using BD Cytofix/Cytoperm™; and intracellular staining with mouse anti-human IFN-γ-V450 (B27), TNF-α-PE-Cy7 (MAb11), IL-2-BV650 (5344.111), (all from BD Biosciences, San Jose, CA). Data were acquired using a BD LRSII flow cytometer equipped with BD FACSDiva software (BD Biosciences, San Jose, CA) and analyzed using FlowJo v.10.7 (FlowJo, LLC, Ashland, OR). Gating strategy consisted in selecting lymphocytes and single cells, followed by discarding cells in the dump channel (dead, CD14+, CD19+ and CD56+ cells), and selecting CD3+ cells in the resulting population. Polyfunctional analysis was performed in FlowJo, defining Boolean combinatorial gates for all the markers in the CD3+/CD4+/CD8-population. These results were visualized in SPICE software v6.0 (Roederer et al., 2011). t-SNE analysis was done in concatenated samples (control, SARS-CoV-2, peptide 163 and peptide 164) from 3 donors using the available plugin in FlowJo.

### Partially-match HLA cell lines

EBV-transformed LG2 cell line (10984, IPD-IMGT/HLA), 9068 cell line (BM9, IHWG), and mouse DP4-transfected MN605 cell line (M12C3-DPA1*0103/DPB1*0401; (Williams et al., 2018); kindly provided by Dr. S. Kent, UMMS), were maintained in RPMI 1640 medium supplemented with L-glutamine (2 mM), penicillin (100 U/mL), streptomycin (100 μg/mL) and 10% FBS at 37°C/5% CO_2_.

### Isolation of T cell clones

T cell clones were isolated by limiting dilution (~1 cell per well) using as feeder cells a pool of irradiated heterologous PBMCs in CRPMI medium supplemented with PHA-P (Gibco, Grand Island, NY) at 1:500 and 100 U/mL IL-2. After 12-14 days incubation wells with cell growth were screened for responses to peptides 163/164, 190/191 and 198/199 by IFN-γ ELISA assay. (Invitrogen, San Diego, CA) and following manufacturer’s protocol. Absorbance at 450 nm was acquired in a BMG plate POLARstar Optima plate reader (Offenburg, Germany). Positive responses were assessed using a cutoff value of 2-times over background + 3-times the standard deviation of background. Sixty-seven T cell clones with the highest specific signal were selected for further analysis.

### T cell clones stimulation and blocking assays

T cell clones (5×10^4^ cells per well) were incubated with equal number of irradiated partially-match HLA cell lines pulsed with peptides (1 μg/mL) or DMSO control in CRPMI+10% FBS; supernatants were collected after 24 hours. Duplicated wells for antigens and 6 wells for negative controls were used. Secreted IFN-γ was measured using ELISA assay as described above. For blocking of antigen-stimulation assays, in-house produced antibodies to HLA-DR (LB3.1), HLA-DQ (SPVL-3), HLA-DP (B7/21), or HLA-ABC (w6/32), were added at a final concentration of 10 μg/mL.

### Peptide binding assay

A fluorescence polarization (FP) competition binding assay similar to published ones (Jurewicz et al., 2019; Yin and Stern, 2014) was used to measure spike peptide binding. Soluble DP4 (HLA-DPA1*01:03/DPB1*04:01) with a covalently-linked Clip peptide was prepared essentially as described (Willis et al., 2021), as were DQ5-Clip (HLA-DQA*01:01/DQB1*05:01) (Jiang et al., 2019) and peptide-exchange catalyst HLA-DM (Busch et al., 1998). Human oxytocinase EKKYFAATQFEPLAARL and influenza nucleoprotein AAHSKAFEDLRVSSY peptides were labeled with Alexa Fluor 488 (Alexa_488_) tetrafluorophenyl ester (Invitrogen, Carlsbad, CA) and used as probe peptides for DP4 and DQ5 binding. Binding reactions were carried out at 37°C in 100 mM sodium citrate, 50 mM sodium chloride, 0.1% octyl β-D-glucopyranoside, 5 mM ethylenediaminetetraacetic acid, 0.1% sodium azide, 0.2 mM iodoacetic acid, 1 mM dithio-threitol as described, but with 1 U/μg thrombin added to cleave the Clip linker. Thrombin was inactivated after 3 hrs of reaction using 0.1 mM phenylmethanesulfonyl fluoride, and the reaction was continued for 21 hours before FP measurement using a Victor X5 Multilabel plate reader (PerkinElmer, Shelton, CT). DP4-Clip (500 nM) and DQ5-Clip (300 nM) concentrations were selected to provide 50% maxiumum binding of 25 nM probe peptide in the presence of 500 nM DM. Binding reactions also contained serial dilutions of test peptides with 5-fold dilutions. IC_50_ values were determined as described (Yin and Stern, 2014).

### Tetramer staining

DP4-163/164 PE and APC tetramers were obtained from the NIH Tetramer Core Facility (Emory University, Atlanta, GA). Cells were collected, washed, and stained using a standard protocol which included: staining for dead cells with Live/Dead Fixable Aqua Dead Cell Stain Kit™ (Life Technologies, Thermo Fisher Scientific, Waltham, MA); blocking of Fc receptors with human Ig (Sigma-Aldrich, St. Louis, MO); staining with the mix of DP4-PE and APC tetramers (final concentration of 2-4 μg/mL each) at 37°C for 2 hours; surface staining antibodies CD3-APC-H7, CD4-PerCP-Cy5.5, CD8-APC-R700, CD14-BV510, CD19-BV510, CD56-BV510 were added for the last 20 minutes of incubation, followed by washes and resuspension in buffer for data acquisition. Data was acquired using a BD LRSII flow cytometer equipped with BD FACSDiva software (BD Biosciences, San Jose, CA) and analyzed using FlowJo v. 10.7 (FlowJo, LLC, Ashland, OR). Gating strategy consisted in selecting lymphocytes and single cells, followed by discarding cells in the dump channel (dead, CD14+, CD19+ and CD56+ cells), CD3+/CD4+ cells and assessing the double-staining PE/APC in this population.

### Sorting of DP4-163/164 cells

For tetramer-sorting, T cells were expanded in vitro with peptides 163/164. After 2 weeks, cells were collected, washed and stained using the procedure described before. Cell populations in the PE+/APC+ gate were sorted using a BD FACS Aria Cell Sorter at The University of Massachusetts Medical School Flow Cytometry Core Facility. Sorted cells were washed and frozen at −80°C until use.

### TCR sequencing

RNA was prepared from HCoV S pool-expanded lines, T cell clones, or sorted cells (0.5-1×10^6^ cells) using RNeasy Mini or RNeasy Micro kits from (QIAGEN, Germantown, MD), following user’s manual instructions. Cells were usually frozen in RLT buffer at time of collection. RNA quality and concentration were assessed using the Fragment Analyzer Service at The University of Massachusetts Molecular Biology Core Labs. RNA with RQN above 7.2 were used for sequencing. We used an in-house RACE (Rapid Amplification of cDNA Ends) approach with template-switch effect, adapted from Turchaninova et al. (Turchaninova et al., 2016) to select and amplify human TCRA and TCRB with RT-primers to the constant region of each chain, and a template-switch primer used introduce unique molecular identifiers (UMI), for error correction during data processing, as well as TrueSeq R1 sequence. Reverse transcription was performed using ~100 ng of RNA and 1 μM RT-primers (mix of primers recognizing the constant region of TCRA or TCRB; Integrated DNA Technologies, IDT, Coralville, IA), and annealed for 3 minutes at 72°C. A reaction mix was added to a final concentration of 1 μM UMI/R1 oligo (IDT), 5 U/μL SMARTScribe reverse transcriptase, 0.5 mM dNTP, 2 U/μL RNAse inhibitor (all Takara Bio USA, Inc, Mountain View, CA), 5 mM DTT (Invitrogen), 1 M betaine (Affymetrix), 6 mM MgCl_2_ (Invitrogen, Thermo Fisher Scientific). Samples were incubated at 42°C for 90 minutes followed by 10 cycles of 50°C/42°C for 2 minutes each, with final incubation at 70°C for 15 minutes. Excess oligo was removed by incubating at 37°C for 40 minutes with 214 U/mL Uracil DNA glycosylase (New England Biolabs, Ipswich, MA). cDNA was purified using AMPure XP Beads (Beckman Coulter, Brea, CA) following manufacturer’s instructions. Four PCR reactions were performed to add TrueSeq R2, P5, and P7 sequences, and i7 indices. All reactions were performed at a final concentration of 0.2 μM primers (IDT), 0.02 U/μl KOD Hot Start DNA Polymerase, 0.2 mM dNTP, 1.5 mM MgSO_4_ (all Novagen / Millipore Sigma, Burlington, MA). All primers sequences shown in STAR Methods. First PCR utilizes purified cDNA, and 2^nd^ strand and RT8 primers; second PCR utilizes purified product from previous PCR, and 2nd strand and nested primers; third PCR utilizes purified product from 2nd PCR, and 5’RACE and barcodes with i7 index primers; fourth PCR utilizes purified product from 3^rd^ PCR, and P1 and P2 primers. Cycling conditions for PCR1: 95°C for 2 minutes; 10 cycles of 95°C for 20 seconds, 70°C for 10 seconds (−1°C per cycle), 70°C for 30 seconds; 15 cycles of 95°C for 20 seconds, 60°C for 10 seconds, 70°C for 30 seconds; final extension at 70°C for 3.5 minutes. PCR2-3: 95°C for 2 minutes; 8 cycles of 95°C for 20 seconds, 60°C for 10 seconds, 70°C for 30 seconds; final extension at 70°C for 3.5 minutes. PCR4: 95°C for 2 minutes; 7 cycles of 95°C for 20 seconds, 60°C for 10 seconds, 70°C for 30 seconds; final extension at 70°C for 3.5 minutes. PCR products from PCR1-3 were purified using AMPure XP magnetics beads, and final PCR product was purified using QIAquick Gel Extraction kit (QIAGEN, Germantown, MD); TCRA and TCRB libraries were quantified using the Fragment Analyzer Service. TCR Sequencing was performed at The University of Massachusetts Deep Sequencing Core. Equimolar concentrations of 8-12 libraries were mixed per lane and sequenced in an Illumina MiSeq System, setup for 250×250 paired end reads. Data was de-multiplexed and single FASTQ files generated for each sample. These files were processed using MIGEC v1.2.9 pipeline: Checkout-batch for de-multiplexing and UMI tag extraction, Histogram for MIG (molecular identifier groups) statistics, and Assemble-batch to perform UMI-guided assembly (Shugay et al., 2014); followed by MiXCR v3.0.13: analyze amplicon pipeline, to align, assemble and export clonotypes (Bolotin et al., 2015). Further data analysis included VDJTools, for gene usage and statistics (Shugay et al., 2014); TCRDist, for clustering trees and logos (Dash et al., 2017); and GLIPH, to find very similar TCRs or TCR with shared motifs (Glanville et al., 2017).

### Clonotype analyses

T cell lines expanded from donors by a single in vitro stimulation with HCoV S pools will likely contain both cross-reactive and HCoV-specific populations, as well as non-specific TCRs. The peptide-expanded lines studied here (Supplemental Tables S5 and S6) showed a skewed distribution of clonotype frequencies relative to a set of TCRβ repertoires defined for samples of unstimulated total PBMC or CD4 cells from non-exposed donors, indicative of a broad expansion of many clonotypes with a range of relative abundances. Analysis of the frequency of CDR3 sequences in the expanded T cell lines that are also observed in DP4-163/164-tetramer sorted cells from other donors, as compared to their frequency in the unexpanded PBMC and CD4 samples, reveals a 7.5-fold greater abundance (p=0.03), allowing us to estimate an approximate false-discovery rate in the expanded polyclonal lines of ~10-20%. The T cell clones were generated by limiting dilution into wells containing irradiated allogenic feeder PBMCs, and sequencing these preparations revealed multiple TCRα and TCRβ sequences in all samples, usually with one or more dominant TCRα and TCRβ clonotypes. Some of the CDR3 sequences were observed in multiple clones, and some clones shared both TCRα and TCRβ sequences. Using this information, we defined candidate TCRα/TCRβ pairs for DQ5 and DP4, paired by detection in multiple clones. In addition, candidate TCRα/TCRβ pairs for DP2 and DP4 were defined, in these cases with pairing assumed from abundance, but not confirmed in independent clones.

### Peptides

Peptides for these studies were obtained from 21st Century Biochemicals (Marlborough, MA), BEI Resources (Manassas, VA), and JPT (Berlin, Germany). Peptide sequences are shown in Supplemental Table S4. HLA-peptide binding prediction was performed with NetMHCIIpan v4.0 server (Reynisson et al., 2020).

## QUANTIFICATION AND STATISTICAL ANALYSIS

Statistical analyses were performed in GraphPad Prism v9.2.0. Comparison between groups were done using Mann-Whitney tests or paired t-tests. ELISpot statistical analysis was performed using a distribution-free resampling (DFR) algorithm described by Moodie et al. (Moodie et al., 2012), and available as online resource at https://rundfr.fredhutch.org.

## List of Supplemental information

Supplemental Table S1. Donors used in the study

Supplemental Table S2. Summary of T cell clones responding to peptides 163/164

Supplemental Table S3. Predicted T cell epitopes in peptides 163/164, 190/191, and 198/199, for MHC-II alleles expressed by donors d0801, d1102, d1202

Supplemental Table S4. Sequences of peptides used in this work

Supplemental Table S5. TCRA and TCRB clonotypes identified in DP4/164-sorted samples.

Supplemental Table S6. TCRA and TCRB clonotypes in HCoV-expanded lines from COVID-19 (d0801, d1102, d1202, G06, G11, G18) and a prepandemic donor (L38).

Supplemental Table S7. Shared CDR3 sequence (identical sequences) in DP4-sorted cells, T cell clones and expanded polyclonal lines.

Supplemental Table S8. Shared CDR3 sequence (identical) with data reported by others: (a) Nolan et al. (2020); (b) Low et al. (2021); (c) Dykema et al. (2021).

Supplemental Table S9. TCR convergence groups (GLIPH)

Supplemental Fig. S1. Functional characterization of in-vitro-expanded cross-reactive cells responding to 190/191 and 198/199. A. Representative intracellular staining plots for IFN-γ, TNF-α, and IL-2 production, and CD107a mobilization to surface in the CD3+ population after re-stimulation of cross-reactive in-vitro expanded T cells with SARS-CoV-2 S pool (CoV-2 S) or individual peptides 190 and 191 (donor d0801), and 198 and 199 (donor d1102). Positive responses shown in red boxes (>3-fold background response). B. Visualization of the polyfunctional response using SPICE (Roederer et al., 2011): pie and arcs graphs showing the combined contribution of each marker.

Supplemental Fig. S2. Sequences, peptide binding motifs, and allelic frequencies for DP4, DP2, and DP402. A. Systematic nomenclature for DPα and DPβ subunits of DP4, DP2, and DP402 allelic variants. All carry the same DPα subunit. B. Peptide binding motifs are similar for all three DP proteins. From NetMHCIIpan4.0. C. DPβ subunit sequences, differences from DP4 are indicated in magenta. From IMGT/HLA database D. Locations of allelic differences on shown on DP2 structure (from PDB:3QLZ, (Dai et al., 2010)). Self-peptide bound to DP2 shown in yellow, with DP4 and DP402 sequence differences shown. Right, schematic diagram of variant DP residues relative to major peptide side-chain binding pockets P1, P4, P6, P7, and P9. Peptide positions P2, P5, and P8 are oriented towards TCR. E. Frequency in various geographic areas of DP4, DP2, and DP402 in various populations, with combined frequency of at least one of these alleles (DP4/2/402). DQ5 frequencies shown for comparison. From IEDB allele frequency tool used to display data in HLA allele frequency database.

Supplemental Fig. S3. HLA-DQ5 and HLA-DP4 competition binding assays for 163/164. Binding of peptide 163/164 (S811-831) to recombinant proteins DP4 (DPA1*01:03/DPB1*04:01) and DQ5 (DQA1*01:01/DQB1*05:01), using competitor peptide from human oxytocinase_271-287_ for DP4 and Influenza A NP peptide for DQ5. IC_50_ for each peptide in each assay shown in parenthesis).

Supplemental Fig. S4. Additional DPA1*01:03/DPB1*04:01-163/164 tetramer staining. A. T cell clones: four T cell clones categorized as DP4-restricted (left), two DP2-restricted (middle), and four DQ5-restricted (right) are shown. B. Ex-vivo staining of unstimulated PBMCs from an unexposed donor. C. Staining of in-vitro HCoV S pool-expanded T cells from pre-pandemic and vaccine donors that expressed DP402 but not DP401. Double-tetramer (PE and APC) staining in CD4+ population is shown in dot plots. DP4-Clip tetramers used as controls. Positive responses shown in red boxes (>3-fold background response).

Supplemental Fig. S5. Cross-reactive response of 163/164-expanded T cell lines from an uninfected donor. A. DPA1*01:03/DPB1*04:01-163/164 tetramer staining of cells expanded with SARS-CoV-2 163/164, HKU1-151, OC43-151, NL63-146, or 229E-115. Double-tetramer (PE and APC) staining in CD4+ population is shown in dot plots; DP4-Clip tetramers used as controls. Positive responses shown in red boxes (>3-fold background response). B. IFN-γ ELISpot of same lines responding to the peptide used for expansion (Expanding peptide) and the cross-reactive peptide(s) (Test peptide) in two uninfected donors. DMSO is the negative control.

## References

Bacher, P., Rosati, E., Esser, D., Martini, G.R., Saggau, C., Schiminsky, E., Dargvainiene, J., Schroder, I., Wieters, I., Khodamoradi, Y., et al. (2020). Low-Avidity CD4(+) T Cell Responses to SARS-CoV-2 in Unexposed Individuals and Humans with Severe COVID-19. Immunity 53, 1258–1271 e1255. 10.1016/j.immuni.2020.11.016

Becerra-Artiles, A., Cruz, J., Leszyk, J.D., Sidney, J., Sette, A., Shaffer, S.A., and Stern, L.J. (2019). Naturally processed HLA-DR3-restricted HHV-6B peptides are recognized broadly with polyfunctional and cytotoxic CD4 T-cell responses. Eur J Immunol 49, 1167–1185. 10.1002/eji.201948126

Bilich, T., Nelde, A., Heitmann, J.S., Maringer, Y., Roerden, M., Bauer, J., Rieth, J., Wacker, M., Peter, A., Horber, S., et al. (2021). T cell and antibody kinetics delineate SARS-CoV-2 peptides mediating long-term immune responses in COVID-19 convalescent individuals. Sci Transl Med 13. 10.1126/scitranslmed.abf7517

Bolotin, D.A., Poslavsky, S., Mitrophanov, I., Shugay, M., Mamedov, I.Z., Putintseva, E.V., and Chudakov, D.M. (2015). MiXCR: software for comprehensive adaptive immunity profiling. Nat Methods 12, 380–381. 10.1038/nmeth.3364

Bonifacius, A., Tischer-Zimmermann, S., Dragon, A.C., Gussarow, D., Vogel, A., Krettek, U., Godecke, N., Yilmaz, M., Kraft, A.R.M., Hoeper, M.M., et al. (2021). COVID-19 immune signatures reveal stable antiviral T cell function despite declining humoral responses. Immunity 54, 340–354 e346. 10.1016/j.immuni.2021.01.008

Braun, J., Loyal, L., Frentsch, M., Wendisch, D., Georg, P., Kurth, F., Hippenstiel, S., Dingeldey, M., Kruse, B., Fauchere, F., et al. (2020). SARS-CoV-2-reactive T cells in healthy donors and patients with COVID-19. Nature 587, 270–274. 10.1038/s41586-020-2598-9

Busch, R., Doebele, R.C., von Scheven, E., Fahrni, J., and Mellins, E.D. (1998). Aberrant intermolecular disulfide bonding in a mutant HLA-DM molecule: implications for assembly, maturation, and function. J Immunol 160, 734–743

Castelli, F.A., Buhot, C., Sanson, A., Zarour, H., Pouvelle-Moratille, S., Nonn, C., Gahery-Segard, H., Guillet, J.G., Menez, A., Georges, B., et al. (2002). HLA-DP4, the most frequent HLA II molecule, defines a new supertype of peptide-binding specificity. J Immunol 169, 6928–6934. 10.4049/jimmunol.169.12.6928

Chen, H., Hou, J., Jiang, X., Ma, S., Meng, M., Wang, B., Zhang, M., Zhang, M., Tang, X., Zhang, F., et al. (2005). Response of memory CD8+ T cells to severe acute respiratory syndrome (SARS) coronavirus in recovered SARS patients and healthy individuals. J Immunol 175, 591–598. 10.4049/jimmunol.175.1.591

Chen, J., Qi, T., Liu, L., Ling, Y., Qian, Z., Li, T., Li, F., Xu, Q., Zhang, Y., Xu, S., et al. (2020). Clinical progression of patients with COVID-19 in Shanghai, China. J Infect 80, e1–e6. 10.1016/j.jinf.2020.03.004

Chen, Z., Boon, S.S., Wang, M.H., Chan, R.W.Y., and Chan, P.K.S. (2021). Genomic and evolutionary comparison between SARS-CoV-2 and other human coronaviruses. J Virol Methods 289, 114032.10.1016/j.jviromet.2020.114032

Cui, J., Li, F., and Shi, Z.L. (2019). Origin and evolution of pathogenic coronaviruses. Nat Rev Microbiol 17, 181–192. 10.1038/s41579-018-0118-9

Dash, P., Fiore-Gartland, A.J., Hertz, T., Wang, G.C., Sharma, S., Souquette, A., Crawford, J.C., Clemens, E.B., Nguyen, T.H.O., Kedzierska, K., et al. (2017). Quantifiable predictive features define epitope-specific T cell receptor repertoires. Nature 547, 89–93. 10.1038/nature22383

Deng, J., Pan, J., Qiu, M., Mao, L., Wang, Z., Zhu, G., Gao, L., Su, J., Hu, Y., Luo, O.J., et al. (2021). Identification of HLA-A2 restricted CD8(+) T cell epitopes in SARS-CoV-2 structural proteins. J Leukoc Biol 110, 1171–1180. 10.1002/JLB.4MA0621-020R

Di Mitri, D., Azevedo, R.I., Henson, S.M., Libri, V., Riddell, N.E., Macaulay, R., Kipling, D., Soares, M.V., Battistini, L., and Akbar, A.N. (2011). Reversible senescence in human CD4+CD45RA+CD27-memory T cells. J Immunol 187, 2093–2100. 10.4049/jimmunol.1100978

Du, R.H., Liang, L.R., Yang, C.Q., Wang, W., Cao, T.Z., Li, M., Guo, G.Y., Du, J., Zheng, C.L., Zhu, Q., et al. (2020). Predictors of mortality for patients with COVID-19 pneumonia caused by SARS-CoV-2: a prospective cohort study. Eur Respir J 55. 10.1183/13993003.00524-2020

Dykema, A.G., Zhang, B., Woldemeskel, B.A., Garliss, C.C., Cheung, L.S., Choudhury, D., Zhang, J., Aparicio, L., Bom, S., Rashid, R., et al. (2021). Functional characterization of CD4+ T cell receptors crossreactive for SARS-CoV-2 and endemic coronaviruses. J Clin Invest 131. 10.1172/JCI146922

Edridge, A.W.D., Kaczorowska, J., Hoste, A.C.R., Bakker, M., Klein, M., Loens, K., Jebbink, M.F., Matser, A., Kinsella, C.M., Rueda, P., et al. (2020). Seasonal coronavirus protective immunity is short-lasting. Nat Med 26, 1691–1693. 10.1038/s41591-020-1083-1

Forni, D., Cagliani, R., Clerici, M., and Sironi, M. (2017). Molecular Evolution of Human Coronavirus Genomes. Trends Microbiol 25, 35–48. 10.1016/j.tim.2016.09.001

Gaunt, E.R., Hardie, A., Claas, E.C., Simmonds, P., and Templeton, K.E. (2010). Epidemiology and clinical presentations of the four human coronaviruses 229E, HKU1, NL63, and OC43 detected over 3 years using a novel multiplex real-time PCR method. J Clin Microbiol 48, 2940–2947. 10.1128/JCM.00636-10

Gerges Harb, J., Noureldine, H.A., Chedid, G., Eldine, M.N., Abdallah, D.A., Chedid, N.F., and Nour-Eldine, W. (2020). SARS, MERS and COVID-19: clinical manifestations and organ-system complications: a mini review. Pathog Dis 78. 10.1093/femspd/ftaa033

Gioia, C., Horejsh, D., Agrati, C., Martini, F., Capobianchi, M.R., Ippolito, G., and Poccia, F. (2005). T-Cell response profiling to biological threat agents including the SARS coronavirus. Int J Immunopathol Pharmacol 18, 525–530. 10.1177/039463200501800312

Glanville, J., Huang, H., Nau, A., Hatton, O., Wagar, L.E., Rubelt, F., Ji, X., Han, A., Krams, S.M., Pettus, C., et al. (2017). Identifying specificity groups in the T cell receptor repertoire. Nature 547, 94–98. 10.1038/nature22976

Gombar, S., Bergquist, T., Pejaver, V., Hammarlund, N.E., Murugesan, K., Mooney, S., Shah, N., Pinsky, B.A., and Banaei, N. (2021). SARS-CoV-2 infection and COVID-19 severity in individuals with prior seasonal coronavirus infection. Diagn Microbiol Infect Dis 100, 115338. 10.1016/j.diagmicrobio.2021.115338

Grifoni, A., Sidney, J., Vita, R., Peters, B., Crotty, S., Weiskopf, D., and Sette, A. (2021). SARS-CoV-2 human T cell epitopes: Adaptive immune response against COVID-19. Cell Host Microbe 29, 1076–1092. 10.1016/j.chom.2021.05.010

Grifoni, A., Weiskopf, D., Ramirez, S.I., Mateus, J., Dan, J.M., Moderbacher, C.R., Rawlings, S.A., Sutherland, A., Premkumar, L., Jadi, R.S., et al. (2020). Targets of T Cell Responses to SARS-CoV-2 Coronavirus in Humans with COVID-19 Disease and Unexposed Individuals. Cell 181, 1489–1501 e1415. 10.1016/j.cell.2020.05.015

Jiang, W., Birtley, J.R., Hung, S.C., Wang, W., Chiou, S.H., Macaubas, C., Kornum, B., Tian, L., Huang, H., Adler, L., et al. (2019). In vivo clonal expansion and phenotypes of hypocretin-specific CD4(+) T cells in narcolepsy patients and controls. Nat Commun 10, 5247. 10.1038/s41467-019-13234-x

Jurewicz, M.M., Willis, R.A., Ramachandiran, V., Altman, J.D., and Stern, L.J. (2019). MHC-I peptide binding activity assessed by exchange after cleavage of peptide covalently linked to beta2-microglobulin. Anal Biochem 584, 113328. 10.1016/j.ab.2019.05.017

Knierman, M.D., Lannan, M.B., Spindler, L.J., McMillian, C.L., Konrad, R.J., and Siegel, R.W. (2020). The Human Leukocyte Antigen Class II Immunopeptidome of the SARS-CoV-2 Spike Glycoprotein. Cell Rep 33, 108454. 10.1016/j.celrep.2020.108454

Koch, C.M., Prigge, A.D., Anekalla, K.R., Shukla, A., Do Umehara, H.C., Setar, L., Chavez, J., Abdala-Valencia, H., Politanska, Y., Markov, N.S., et al. (2021). Age-related Differences in the Nasal Mucosal Immune Response to SARS-CoV-2. Am J Respir Cell Mol Biol. 10.1165/rcmb.2021-0292OC

Le Bert, N., Tan, A.T., Kunasegaran, K., Tham, C.Y.L., Hafezi, M., Chia, A., Chng, M.H.Y., Lin, M., Tan, N., Linster, M., et al. (2020). SARS-CoV-2-specific T cell immunity in cases of COVID-19 and SARS, and uninfected controls. Nature 584, 457–462. 10.1038/s41586-020-2550-z

Lipsitch, M., Grad, Y.H., Sette, A., and Crotty, S. (2020). Cross-reactive memory T cells and herd immunity to SARS-CoV-2. Nat Rev Immunol 20, 709–713. 10.1038/s41577-020-00460-4

Liu, W., Fontanet, A., Zhang, P.H., Zhan, L., Xin, Z.T., Baril, L., Tang, F., Lv, H., and Cao, W.C. (2006). Two-year prospective study of the humoral immune response of patients with severe acute respiratory syndrome. J Infect Dis 193, 792–795. 10.1086/500469

Low, J.S., Vaqueirinho, D., Mele, F., Foglierini, M., Jerak, J., Perotti, M., Jarrossay, D., Jovic, S., Perez, L., Cacciatore, R., et al. (2021). Clonal analysis of immunodominance and cross-reactivity of the CD4 T cell response to SARS-CoV-2. Science 372, 1336–1341. 10.1126/science.abg8985

Loyal, L., Braun, J., Henze, L., Kruse, B., Dingeldey, M., Reimer, U., Kern, F., Schwarz, T., Mangold, M., Unger, C., et al. (2021). Cross-reactive CD4(+) T cells enhance SARS-CoV-2 immune responses upon infection and vaccination. Science 374, eabh1823. 10.1126/science.abh1823

Lu, X., Hosono, Y., Nagae, M., Ishizuka, S., Ishikawa, E., Motooka, D., Ozaki, Y., Sax, N., Maeda, Y., Kato, Y., et al. (2021). Identification of conserved SARS-CoV-2 spike epitopes that expand public cTfh clonotypes in mild COVID-19 patients. J Exp Med 218. 10.1084/jem.20211327

Mateus, J., Grifoni, A., Tarke, A., Sidney, J., Ramirez, S.I., Dan, J.M., Burger, Z.C., Rawlings, S.A., Smith, D.M., Phillips, E., et al. (2020). Selective and cross-reactive SARS-CoV-2 T cell epitopes in unexposed humans. Science 370, 89–94. 10.1126/science.abd3871

Mazzoni, A., Salvati, L., Maggi, L., Annunziato, F., and Cosmi, L. (2021). Hallmarks of immune response in COVID-19: Exploring dysregulation and exhaustion. Semin Immunol, 101508. 10.1016/j.smim.2021.101508

Meyer, B., Drosten, C., and Muller, M.A. (2014). Serological assays for emerging coronaviruses: challenges and pitfalls. Virus Res 194, 175–183. 10.1016/j.virusres.2014.03.018

Moodie, Z., Price, L., Janetzki, S., and Britten, C.M. (2012). Response determination criteria for ELISPOT: toward a standard that can be applied across laboratories. Methods Mol Biol 792, 185–196. 10.1007/978-1-61779-325-7_15

Nelde, A., Bilich, T., Heitmann, J.S., Maringer, Y., Salih, H.R., Roerden, M., Lubke, M., Bauer, J., Rieth, J., Wacker, M., et al. (2021). SARS-CoV-2-derived peptides define heterologous and COVID-19-induced T cell recognition. Nat Immunol 22, 74–85. 10.1038/s41590-020-00808-x

Ng, K.W., Faulkner, N., Cornish, G.H., Rosa, A., Harvey, R., Hussain, S., Ulferts, R., Earl, C., Wrobel, A.G., Benton, D.J., et al. (2020). Preexisting and de novo humoral immunity to SARS-CoV-2 in humans. Science 370, 1339–1343. 10.1126/science.abe1107

Nolan, S., Vignali, M., Klinger, M., Dines, J.N., Kaplan, I.M., Svejnoha, E., Craft, T., Boland, K., Pesesky, M., Gittelman, R.M., et al. (2020). A large-scale database of T-cell receptor beta (TCRbeta) sequences and binding associations from natural and synthetic exposure to SARS-CoV-2. Res Sq. 10.21203/rs.3.rs-51964/v1

Parker, R., Partridge, T., Wormald, C., Kawahara, R., Stalls, V., Aggelakopoulou, M., Parker, J., Powell Doherty, R., Ariosa Morejon, Y., Lee, E., et al. (2021). Mapping the SARS-CoV-2 spike glycoprotein-derived peptidome presented by HLA class II on dendritic cells. Cell Rep 35, 109179. 10.1016/j.celrep.2021.109179

Peng, Y., Mentzer, A.J., Liu, G., Yao, X., Yin, Z., Dong, D., Dejnirattisai, W., Rostron, T., Supasa, P., Liu, C., et al. (2020). Broad and strong memory CD4(+) and CD8(+) T cells induced by SARS-CoV-2 in UK convalescent individuals following COVID-19. Nat Immunol 21, 1336–1345. 10.1038/s41590-020-0782-6

Poh, C.M., Carissimo, G., Wang, B., Amrun, S.N., Lee, C.Y., Chee, R.S., Fong, S.W., Yeo, N.K., Lee, W.H., Torres-Ruesta, A., et al. (2020). Two linear epitopes on the SARS-CoV-2 spike protein that elicit neutralising antibodies in COVID-19 patients. Nat Commun 11, 2806. 10.1038/s41467-020-16638-2

Reynisson, B., Alvarez, B., Paul, S., Peters, B., and Nielsen, M. (2020). NetMHCpan-4.1 and NetMHCIIpan-4.0: improved predictions of MHC antigen presentation by concurrent motif deconvolution and integration of MS MHC eluted ligand data. Nucleic Acids Res 48, W449–W454. 10.1093/nar/gkaa379

Roederer, M., Nozzi, J.L., and Nason, M.C. (2011). SPICE: exploration and analysis of post-cytometric complex multivariate datasets. Cytometry A 79, 167–174. 10.1002/cyto.a.21015

Rossjohn, J., Gras, S., Miles, J.J., Turner, S.J., Godfrey, D.I., and McCluskey, J. (2015). T cell antigen receptor recognition of antigen-presenting molecules. Annu Rev Immunol 33, 169–200. 10.1146/annurevimmunol-032414-112334

Rydyznski Moderbacher, C., Ramirez, S.I., Dan, J.M., Grifoni, A., Hastie, K.M., Weiskopf, D., Belanger, S., Abbott, R.K., Kim, C., Choi, J., et al. (2020). Antigen-Specific Adaptive Immunity to SARS-CoV-2 in Acute COVID-19 and Associations with Age and Disease Severity. Cell 183, 996–1012 e1019. 10.1016/j.cell.2020.09.038

Sagar, M., Reifler, K., Rossi, M., Miller, N.S., Sinha, P., White, L.F., and Mizgerd, J.P. (2021). Recent endemic coronavirus infection is associated with less-severe COVID-19. J Clin Invest 131. 10.1172/JCI143380

Saini, S.K., Hersby, D.S., Tamhane, T., Povlsen, H.R., Amaya Hernandez, S.P., Nielsen, M., Gang, A.O., and Hadrup, S.R. (2021). SARS-CoV-2 genome-wide T cell epitope mapping reveals immunodominance and substantial CD8(+) T cell activation in COVID-19 patients. Sci Immunol 6. 10.1126/sciimmunol.abf7550

Sette, A., and Crotty, S. (2021). Adaptive immunity to SARS-CoV-2 and COVID-19. Cell 184, 861–880. 10.1016/j.cell.2021.01.007

Shrock, E., Fujimura, E., Kula, T., Timms, R.T., Lee, I.H., Leng, Y., Robinson, M.L., Sie, B.M., Li, M.Z., Chen, Y., et al. (2020). Viral epitope profiling of COVID-19 patients reveals cross-reactivity and correlates of severity. Science 370. 10.1126/science.abd4250

Shugay, M., Britanova, O.V., Merzlyak, E.M., Turchaninova, M.A., Mamedov, I.Z., Tuganbaev, T.R., Bolotin, D.A., Staroverov, D.B., Putintseva, E.V., Plevova, K., et al. (2014). Towards error-free profiling of immune repertoires. Nat Methods 11, 653–655. 10.1038/nmeth.2960

Sidney, J., Steen, A., Moore, C., Ngo, S., Chung, J., Peters, B., and Sette, A. (2010a). Divergent motifs but overlapping binding repertoires of six HLA-DQ molecules frequently expressed in the worldwide human population. J Immunol 185, 4189–4198. 10.4049/jimmunol.1001006

Sidney, J., Steen, A., Moore, C., Ngo, S., Chung, J., Peters, B., and Sette, A. (2010b). Five HLA-DP molecules frequently expressed in the worldwide human population share a common HLA supertypic binding specificity. J Immunol 184, 2492–2503. 10.4049/jimmunol.0903655

Soveg, F.W., Schwerk, J., Gokhale, N.S., Cerosaletti, K., Smith, J.R., Pairo-Castineira, E., Kell, A.M., Forero, A., Zaver, S.A., Esser-Nobis, K., et al. (2021). Endomembrane targeting of human OAS1 p46 augments antiviral activity. Elife 10. 10.7554/eLife.71047

Stern, L.J., and Wiley, D.C. (1994). Antigenic peptide binding by class I and class II histocompatibility proteins. Structure 2, 245–251. 10.1016/s0969-2126(00)00026-5

Stockton, J.D., Nieto, T., Wroe, E., Poles, A., Inston, N., Briggs, D., and Beggs, A.D. (2020). Rapid, highly accurate and cost-effective open-source simultaneous complete HLA typing and phasing of class I and II alleles using nanopore sequencing. HLA 96, 163–178. 10.1111/tan.13926

Swadling, L., Diniz, M.O., Schmidt, N.M., Amin, O.E., Chandran, A., Shaw, E., Pade, C., Gibbons, J.M., Le Bert, N., Tan, A.T., et al. (2021). Pre-existing polymerase-specific T cells expand in abortive seronegative SARS-CoV-2. Nature. 10.1038/s41586-021-04186-8

Tan, A.T., Linster, M., Tan, C.W., Le Bert, N., Chia, W.N., Kunasegaran, K., Zhuang, Y., Tham, C.Y.L., Chia, A., Smith, G.J.D., et al. (2021a). Early induction of functional SARS-CoV-2-specific T cells associates with rapid viral clearance and mild disease in COVID-19 patients. Cell Rep 34, 108728. 10.1016/j.celrep.2021.108728

Tan, C.C.S., Owen, C.J., Tham, C.Y.L., Bertoletti, A., van Dorp, L., and Balloux, F. (2021b). Pre-existing T cell-mediated cross-reactivity to SARS-CoV-2 cannot solely be explained by prior exposure to endemic human coronaviruses. Infect Genet Evol 95, 105075. 10.1016/j.meegid.2021.105075

Tarke, A., Sidney, J., Kidd, C.K., Dan, J.M., Ramirez, S.I., Yu, E.D., Mateus, J., da Silva Antunes, R., Moore, E., Rubiro, P., et al. (2021). Comprehensive analysis of T cell immunodominance and immunoprevalence of SARS-CoV-2 epitopes in COVID-19 cases. Cell Rep Med 2, 100204. 10.1016/j.xcrm.2021.100204

Tian, Y., Babor, M., Lane, J., Schulten, V., Patil, V.S., Seumois, G., Rosales, S.L., Fu, Z., Picarda, G., Burel, J., et al. (2017). Unique phenotypes and clonal expansions of human CD4 effector memory T cells re-expressing CD45RA. Nat Commun 8, 1473. 10.1038/s41467-017-01728-5

Turchaninova, M.A., Davydov, A., Britanova, O.V., Shugay, M., Bikos, V., Egorov, E.S., Kirgizova, V.I., Merzlyak, E.M., Staroverov, D.B., Bolotin, D.A., et al. (2016). High-quality full-length immunoglobulin profiling with unique molecular barcoding. Nat Protoc 11, 1599–1616. 10.1038/nprot.2016.093

Vanderbeke, L., Van Mol, P., Van Herck, Y., De Smet, F., Humblet-Baron, S., Martinod, K., Antoranz, A., Arijs, I., Boeckx, B., Bosisio, F.M., et al. (2021). Monocyte-driven atypical cytokine storm and aberrant neutrophil activation as key mediators of COVID-19 disease severity. Nat Commun 12, 4117. 10.1038/s41467-021-24360-w

Verhagen, J., van der Meijden, E.D., Lang, V., Kremer, A.E., Volkl, S., Mackensen, A., Aigner, M., and Kremer, A.N. (2021). Human CD4(+) T cells specific for dominant epitopes of SARS-CoV-2 Spike and Nucleocapsid proteins with therapeutic potential. Clin Exp Immunol 205, 363–378. 10.1111/cei.13627

Voss, C., Esmail, S., Liu, X., Knauer, M.J., Ackloo, S., Kaneko, T., Lowes, L., Stogios, P., Seitova, A., Hutchinson, A., et al. (2021). Epitope-specific antibody responses differentiate COVID-19 outcomes and variants of concern. JCI Insight 6. 10.1172/jci.insight.148855

Ward, J.D., Cornaby, C., and Schmitz, J.L. (2021). Indeterminate QuantiFERON Gold Plus Results Reveal Deficient Interferon Gamma Responses in Severely Ill COVID-19 Patients. J Clin Microbiol 59, e0081121. 10.1128/JCM.00811-21

Weiskopf, D., Schmitz, K.S., Raadsen, M.P., Grifoni, A., Okba, N.M.A., Endeman, H., van den Akker, J.P.C., Molenkamp, R., Koopmans, M.P.G., van Gorp, E.C.M., et al. (2020). Phenotype and kinetics of SARS-CoV-2-specific T cells in COVID-19 patients with acute respiratory distress syndrome. Sci Immunol 5. 10.1126/sciimmunol.abd2071

Wickenhagen, A., Sugrue, E., Lytras, S., Kuchi, S., Noerenberg, M., Turnbull, M.L., Loney, C., Herder, V., Allan, J., Jarmson, I., et al. (2021). A prenylated dsRNA sensor protects against severe COVID-19. Science 374, eabj3624. 10.1126/science.abj3624

Wiersinga, W.J., Rhodes, A., Cheng, A.C., Peacock, S.J., and Prescott, H.C. (2020). Pathophysiology, Transmission, Diagnosis, and Treatment of Coronavirus Disease 2019 (COVID-19): A Review. JAMA 324, 782–793. 10.1001/jama.2020.12839

Williams, T., Krovi, H.S., Landry, L.G., Crawford, F., Jin, N., Hohenstein, A., DeNicola, M.E., Michels, A.W., Davidson, H.W., Kent, S.C., et al. (2018). Development of T cell lines sensitive to antigen stimulation. J Immunol Methods 462, 65–73. 10.1016/j.jim.2018.08.011

Willis, R.A., Ramachandiran, V., Shires, J.C., Bai, G., Jeter, K., Bell, D.L., Han, L., Kazarian, T., Ugwu, K.C., Laur, O., et al. (2021). Production of Class II MHC Proteins in Lentiviral Vector-Transduced HEK-293T Cells for Tetramer Staining Reagents. Curr Protoc 1, e36. 10.1002/cpz1.36

Woldemeskel, B.A., Garliss, C.C., and Blankson, J.N. (2021). mRNA Vaccine-Elicited SARS-CoV-2-Specific T cells Persist at 6 Months and Recognize the Delta Variant. Clin Infect Dis. 10.1093/cid/ciab915

Xia, X. (2021). Domains and Functions of Spike Protein in Sars-Cov-2 in the Context of Vaccine Design. Viruses 13. 10.3390/v13010109

Yang, J., James, E., Roti, M., Huston, L., Gebe, J.A., and Kwok, W.W. (2009). Searching immunodominant epitopes prior to epidemic: HLA class II-restricted SARS-CoV spike protein epitopes in unexposed individuals. Int Immunol 21, 63–71. 10.1093/intimm/dxn124

Yin, L., and Stern, L.J. (2014). Measurement of Peptide Binding to MHC Class II Molecules by Fluorescence Polarization. Curr Protoc Immunol 106, 5 10 11–15 10 12. 10.1002/0471142735.im0510s106

Zhao, Y., Kilian, C., Turner, J.E., Bosurgi, L., Roedl, K., Bartsch, P., Gnirck, A.C., Cortesi, F., Schultheiss, C., Hellmig, M., et al. (2021). Clonal expansion and activation of tissue-resident memory-like Th17 cells expressing GM-CSF in the lungs of severe COVID-19 patients. Sci Immunol 6. 10.1126/sciimmunol.abf6692

Ziegler, C.G.K., Miao, V.N., Owings, A.H., Navia, A.W., Tang, Y., Bromley, J.D., Lotfy, P., Sloan, M., Laird, H., Williams, H.B., et al. (2021). Impaired local intrinsic immunity to SARS-CoV-2 infection in severe COVID-19. Cell 184, 4713–4733 e4722. 10.1016/j.cell.2021.07.023

