## Supplemental Figures for "Broadly-recognized, cross-reactive SARS-CoV-2 CD4 T cell epitopes are highly conserved across human coronaviruses and presented by common HLA alleles"

Supplemental Figure S1:

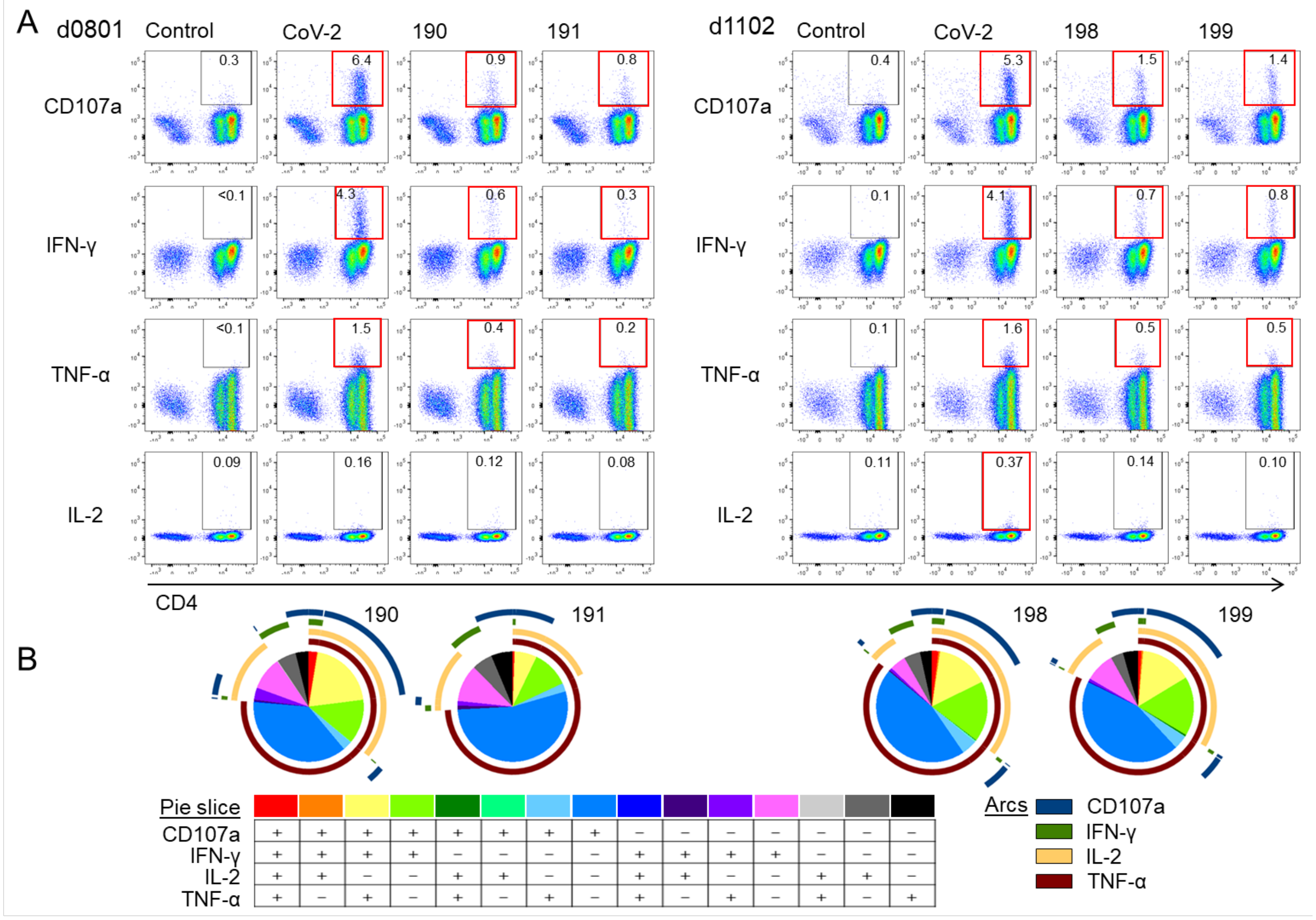


Supplemental Fig. S1. Functional characterization of in-vitro-expanded cross-reactive cells responding to 190/191 and 198/199. A. Representative intracellular staining plots for IFN-γ, TNF-α, and IL-2 production, and CD107a mobilization to surface in the CD3+ population after re-stimulation of cross-reactive in-vitro expanded T cells with SARS-CoV-2 S pool (CoV-2 S) or individual peptides 190 and 191 (donor d0801), and 198 and 199 (donor d1102). Positive responses shown in red boxes (>3-fold background response). B. Visualization of the polyfunctional response using SPICE (Roederer et al., 2011): pie and arcs graphs showing the combined contribution of each marker.

Supplemental Figure S2:

A

DP4: HLA-DPA1*01:03 / HLA-DPB1*04:01
DP2: HLA-DPA1*01:03 / HLA-DPB1*02:01
DP402: HLA-DPA1*01:03 / HLA-DPB1*04:02

B

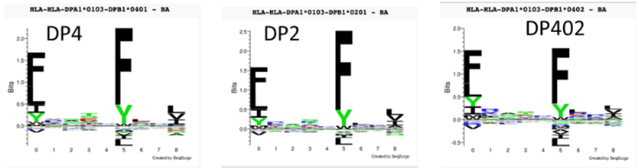


C

```
AA Pos.                     10         20         30         40         50         60         70         80         90        100
DPB1*04:01:01   RATPENYLFQ GRQECYAFNG TQRFLERYIY NREEFARFDS DVGEFRAVTE LGRPAAEYWN SQKDILEEKR AVPDRMCRHN YELGGPMTLQ RRVQPRVNVS
DPB1*02:01:02   ---------- ---------- ---------- -----V---- ---------- ----DE---- --------E- ---------- ---------- ----------
DPB1*04:02:01   ---------- ---------- ---------- -----V---- ---------- ----DE---- ---------- ---------- ---------- ----------

AA Pos.                    110        120        130        140        150        160        170        180        190        200
DPB1*04:01:01   PSKKGPLQHH NLLVCHVTDF YPGSIQVRWF LNGQEETAGV VSTNLIRNGD WTFQILVMLE MTPQQGDVYT CQVEHTSLDS PVTVEWKAQS DSARSKTLTG
DPB1*02:01:02   ---------- ---------- ---------- ---------- ---------- ---------- ---------- ---------- ---------- ----------
DPB1*04:02:01   ---------- ---------- ---------- ---------- ---------- ---------- ---------- -------M-- ---------- ----------

AA Pos.                    210        220
DPB1*04:01:01   AGGFVLGLII CGVGIFMHRR SKKVQRGSA
DPB1*02:01:02   ---------- ---------- ---------
DPB1*04:02:01   ---------- ---------- ---------
```

D

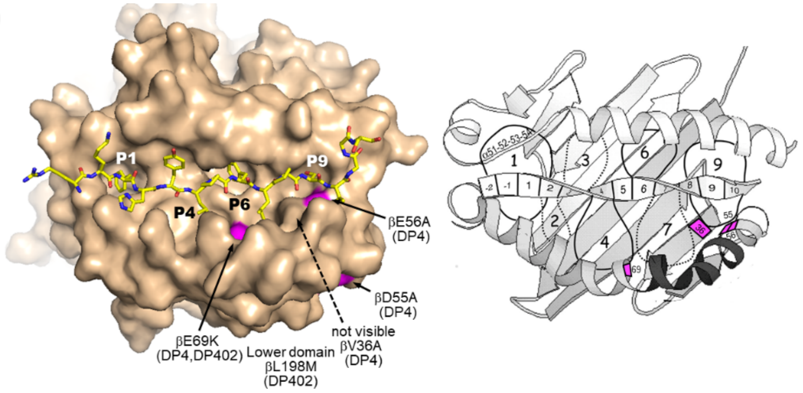


E

| Region | DP4 | DP2 | DP402 | 3DPs | DQ5 |
|---|---|---|---|---|---|
| North America | 37.2 | 17.0 | 45.8 | 71.7 | 2.4 |
| Mexico | 38.1 | 11.3 | 89.5 | 94.2 | 1.7 |
| South America | 12.9 | 6.2 | 36.7 | 48.3 | 1.4 |
| Europe | 55.4 | 26.5 | 18.1 | 73.2 | 3.2 |
| Central Africa | 7.8 | 14.8 | 32.5 | 46.9 | 5.6 |
| East Africa | 4.6 | 10.3 | 19.1 | 30.7 | 12.7 |
| West Africa | 1.5 | 6.7 | 6.9 | 14.4 | 7.7 |
| Northeast Asia | 10.1 | 12.2 | 8.6 | 27.9 | 1.7 |
| East Asia | 1.2 | 3.6 | 1.5 | 6.2 | 2.4 |
| South Asia | 50.1 | 21.5 | 12.5 | 65.7 | 3.8 |
| Oceania | 23.8 | 21.2 | 2.8 | 41.6 | 0.5 |

Supplemental Fig. S2. Sequences, peptide binding motifs, and allelic frequencies for DP4, DP2, and DP402. A. Systematic nomenclature for DPα and DPβ subunits of DP4, DP2, and DP402 allelic variants. All carry the same DPα subunit. B. Peptide binding motifs are similar for all three DP proteins. From NetMHCIIpan4.0. C. DPβ subunit sequences, differences from DP4 are indicated in magenta. From IMGT/HLA database D. Locations of allelic differences on shown on DP2 structure (from PDB:3QLZ, (Dai et al., 2010)). Self-peptide bound to DP2 shown in yellow, with DP4 and DP402 sequence differences shown. Right, schematic diagram of variant DP residues relative to major peptide side-chain binding pockets P1, P4, P6, P7, and P9. Peptide positions P2, P5, and P8 are oriented towards TCR. E. Frequency in various geographic areas of DP4, DP2, and DP402 in various populations, with combined frequency of at least one of these alleles (DP4/2/402). DQ5 frequencies shown for comparison. From IEDB allele frequency tool used to display data in HLA allele frequency database.

Supplemental Figure S3:

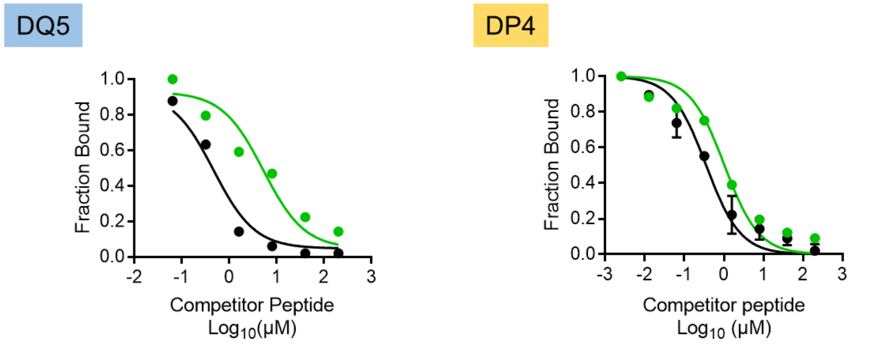


**$S_{811\text{-}831}$:**
KPSKRSFIEDLLFNKVTLADA ($IC_{50}$= 5.24 μM)

**Influenza A nucleoprotein**:
AAHSKAFEDLRVSSY ($IC_{50}$= 0.47 μM)

**$S_{811\text{-}831}$:**
KPSKRSFIEDLLFNKVTLADA ($IC_{50}$= 1.04 μM)

**Human oxytocinase$_{271\text{-}287}$**:
EKKYFAATQFEPLAARL ($IC_{50}$= 0.36 μM)

Supplemental Fig. S3. HLA-DQ5 and HLA-DP4 competition binding assays for 163/164. Binding of peptide 163/164 ($S_{811\text{-}831}$) to recombinant proteins DP4 (DPA1*01:03/DPB1*04:01) and DQ5 (DQA1*01:01/DQB1*05:01), using competitor peptide from human oxytocinase$_{271\text{-}287}$ for DP4 and Influenza A NP peptide for DQ5. $IC_{50}$ for each peptide in each assay shown in parenthesis).

Supplemental Figure S4:

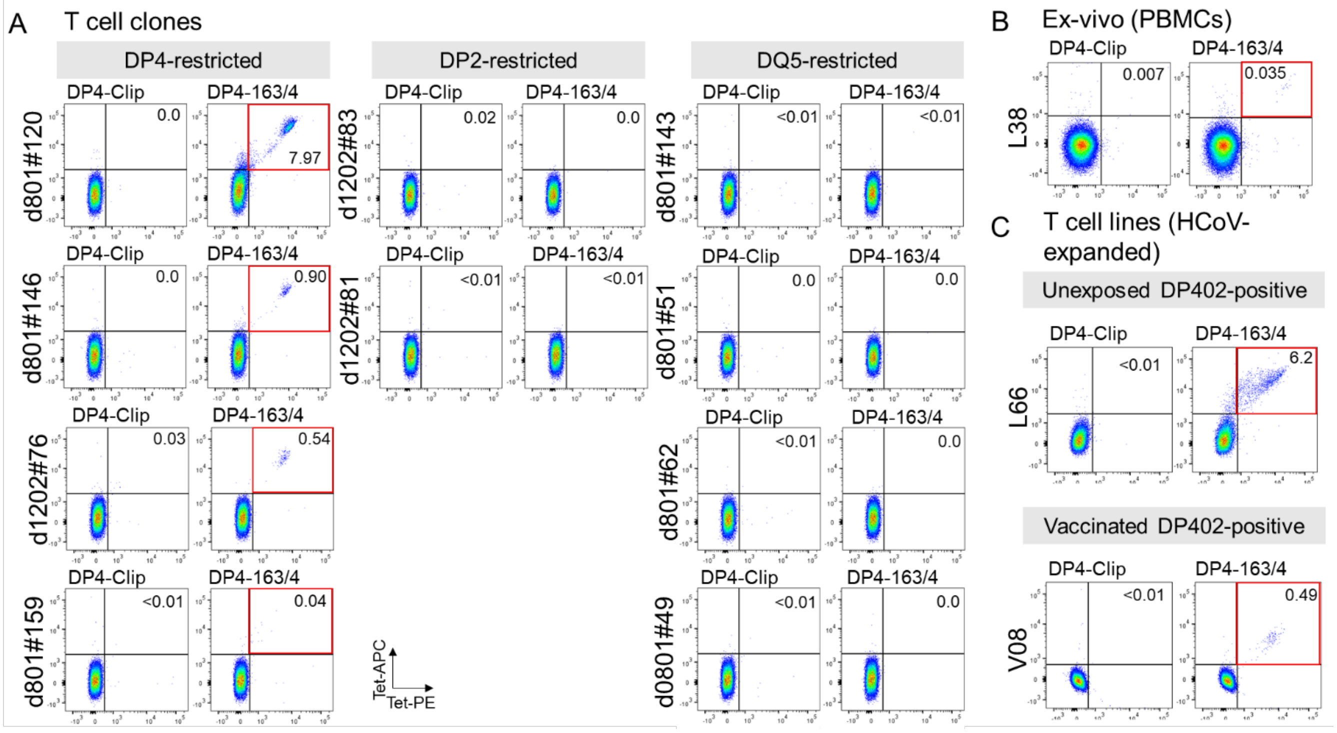


Supplemental Fig. S4. Additional DPA1*01:03/DPB1*04:01-163/164 tetramer staining. A. T cell clones: four T cell clones categorized as DP4-restricted (left), two DP2-restricted (middle), and four DQ5-restricted (right) are shown. B. Ex-vivo staining of unstimulated PBMCs from an unexposed donor. C. Staining of in-vitro HCoV S pool expanded T cells from unexposed, and vaccine donors that expressed DP402 but not DP401. Double-tetramer (PE and APC) staining in CD4+ population is shown in dot plots. DP4-CLIP tetramers used as controls. Positive responses shown in red boxes (>3-fold background response).

Supplemental Figure S5:

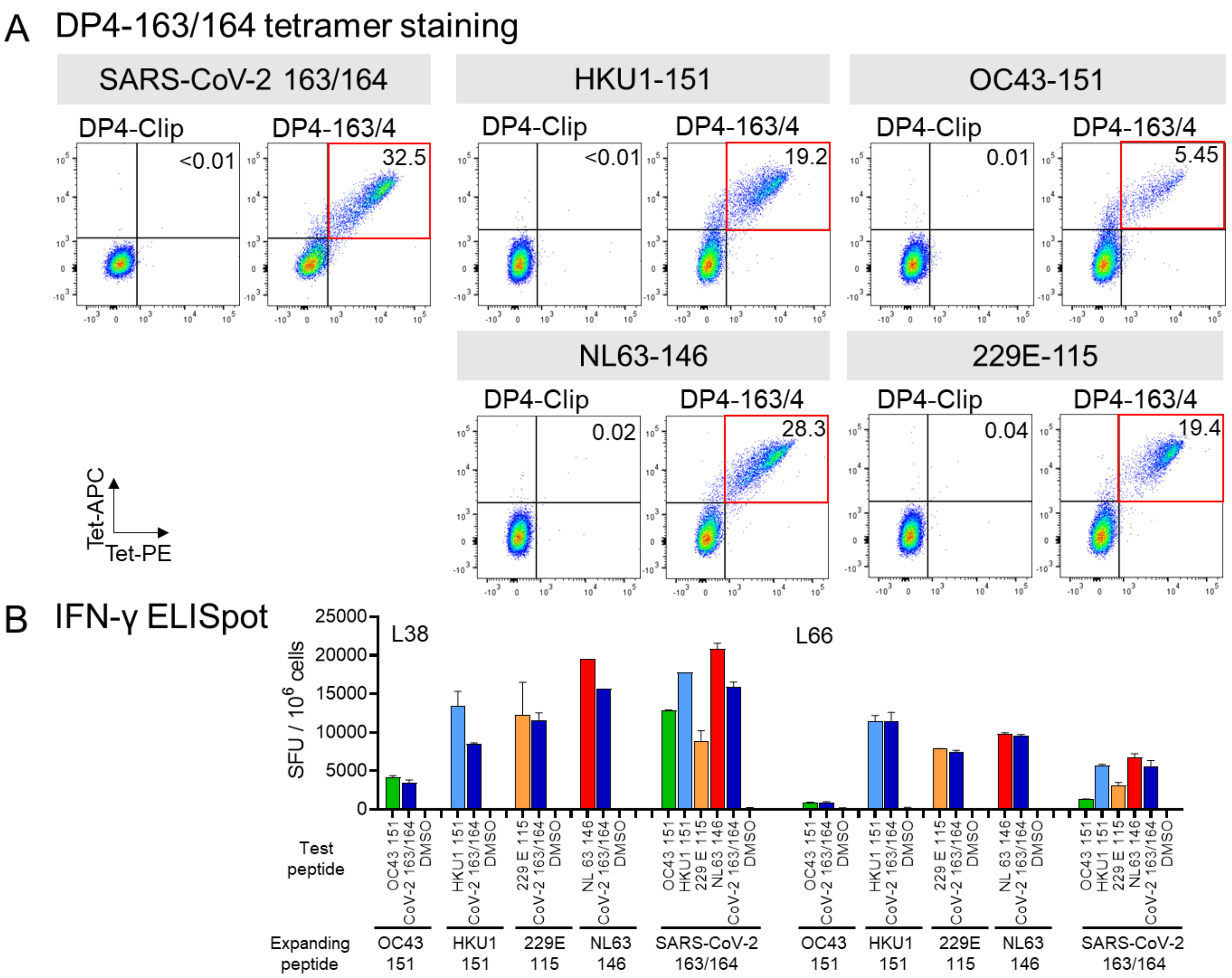


Supplemental Fig. S5. Cross-reactive response of 163/164-expanded T cell lines from an unexposed donor. A. DPA1*01:03/DPB1*04:01-163/164 tetramer staining of cells expanded with SARS-CoV-2 163/164, HKU1-151, OC43-151, NL63-146, or 229E-115. Double-tetramer (PE and APC) staining in CD4+ population is shown in dot plots; DP4-Clip tetramers used as controls. Positive responses shown in red boxes (>3-fold background response). B. IFN-γ ELISpot of same lines responding to the peptide used for expansion (Expanding peptide) and the cross-reactive peptide(s) (Test peptide) in two unexposed donors. DMSO is the negative control.
